# The NUCKS1-SKP2-p21/p27 axis controls S phase entry

**DOI:** 10.1101/2021.06.29.450420

**Authors:** Samuel Hume, Claudia P. Grou, Pauline Lascaux, Vincenzo D’Angiolella, Arnaud J. Legrand, Kristijan Ramadan, Grigory L. Dianov

**Affiliations:** Medical Research Council Oxford Institute for Radiation Oncology, Department of Oncology, University of Oxford, OX3 7DQ, Oxford, UK; Institute of Cytology and Genetics, Siberian Branch of the Russian Academy of Sciences, Lavrentieva 10, 630090 Novosibirsk, Russia; Novosibirsk State University, Novosibirsk, Russian Federation, 630090

## Abstract

Efficient entry into S phase of the cell cycle is necessary for embryonic development and tissue homeostasis. However, unscheduled S phase entry triggers DNA damage and promotes oncogenesis, underlining the requirement for strict control. Here, we identify the NUCKS1-SKP2-p21/p27 axis as a checkpoint pathway for the G1/S transition. In response to mitogenic stimulation, NUCKS1, a transcription factor, is recruited to chromatin to activate expression of *SKP2*, the F-box component of the SCF^SKP2^ ubiquitin ligase, leading to degradation of p21 and p27 and promoting progression into S phase. In contrast, DNA damage induces p53-dependent transcriptional repression of *NUCKS1*, leading to SKP2 downregulation, p21/p27 upregulation, and cell cycle arrest. We propose that the NUCKS1-SKP2-p21/p27 axis integrates mitogenic and DNA damage signalling to control S phase entry. TCGA data reveal that this mechanism is hijacked in many cancers, potentially allowing cancer cells to sustain uncontrolled proliferation.

## Introduction

Entry into S phase of the cell cycle is essential to sustain the proliferation that permits embryonic development and tissue repair^1^, but unscheduled S phase entry induces replication stress, DNA damage, and oncogenesis^2–5^. G1/S progression must therefore be strictly controlled^6–8^. S phase entry is driven by mitogens, which increase the ratio of G1/S cyclins: cyclin-dependent kinase (CDK) inhibitors and activate G1/S CDKs as a result. In contrast, DNA damage inhibits S phase entry, stimulating p53 signalling to reduce the G1/S cyclin: CDK inhibitor ratio and prevent G1/S CDK activity^9^. Only cells whose mitogenic signalling outcompetes their DNA damage load are permitted to enter S phase^6,10–14^, which must be achieved through the integration of these antagonistic stimuli by signalling hubs. However, signalling hubs that achieve this goal are not well-characterised^6^.

The transcription factor Nuclear Ubiquitous Casein kinase and cyclin-dependent Kinase Substrate 1 (NUCKS1) has emerged in the light of recent studies as a promising candidate for one such signalling hub. NUCKS1, a member of the high mobility group family of proteins^15^, increases chromatin accessibility at target promoters to enable the recruitment of RNA polymerase II^16^. So far, the only direct transcriptional targets identified for NUCKS1 regulate insulin receptor signalling^16^. However, NUCKS1 is known to affect cell cycle progression and proliferation in mammary epithelial cells^17^ and gastric cancer cells^18^, and also plays a role in the protection of replication fidelity by regulating double-strand break (DSB) repair^19–22^. In addition, NUCKS1 is a phosphorylation substrate for CDK2 and CDK1, the major kinases controlling the G1/S and G2/M transitions^23–28^, and for the DNA damage response (DDR) kinases ATM and DNA-PK^29,30^. Furthermore, Rb-E2F^31^ and p53^32^ have been detected in the proximity of the *NUCKS1* promoter by genome wide ChIP-Seq, suggesting that *NUCKS1* expression might be regulated by the cell cycle or by DNA damage.

NUCKS1 also exhibits oncogenic properties, and its overexpression, correlating with poor patient prognosis, has been reported in a number of cancers^33–38^. Furthermore, NUCKS1 depletion inhibits - while its overexpression promotes - xenograft tumour growth^18,39,40^, suggesting a direct role in tumourigenesis.

Altogether, these studies suggest a potentially important role for NUCKS1 in cell cycle progression. However, mechanistic details explaining how NUCKS1 does this are unknown. In particular, whether NUCKS1 employs transcriptional control of the cell cycle – and which putative targets of NUCKS1 are involved – has not been established. The precise cell cycle phase affected by NUCKS1 is also not known, and how NUCKS1 is regulated throughout the cell cycle, by mitogens, or following DNA damage, has not been explored.

Here, we show that S phase Kinase-associated Protein 2 (*SKP2*) is a transcriptional target for NUCKS1 in late G1 phase, and identify the SKP2-p21/p27 axis as a pathway controlled by NUCKS1. SKP2 is a substrate-recruiting F-box protein, which forms, along with SKP1, CUL1, and RBX1, the SCF^SKP2^ ubiquitin ligase complex^41^. During the G1/S transition, SKP2 directs SCF^SKP2^ for degradation of the CDK inhibitors p21 and p27, relieving p21/p27-mediated inhibition of cyclin E-CDK2^12,14,42,43^. In this way, SCF^SKP2^ controls cell cycle and cancer progression^44–46^.

We find that the SKP2-p21/p27 axis acts through NUCKS1 to integrate mitogenic and DNA damage signalling at the G1/S transition. We show that NUCKS1 is stimulated by mitogens to promote *SKP2* expression and consequent p21/p27 degradation, enabling S phase entry. In contrast, DNA damage inhibits NUCKS1 through p53, reducing SKP2 levels, increasing p21/p27 levels, and blocking S phase entry. In this way, the NUCKS1-SKP2-p21/p27 axis acts as a checkpoint pathway for the G1/S transition, only permitting S phase entry for cells whose mitogenic signalling outcompetes their load of DNA damage.

## Results

### NUCKS1 transcriptionally controls the SKP2-p21/p27 axis

To investigate whether NUCKS1 regulates the transcription of genes involved in cell cycle progression, we cross-compared a list of genes whose expression correlates with *NUCKS1* mRNA in tumour samples and cell lines^47^, with genes whose promoter NUCKS1 binds in genome-wide ChIP-Seq^16^. This generated a list of 232 putative NUCKS1 target genes. Among them, we found several genes regulating the G1/S transition (e.g., *SKP2, CCND1, CDK6, E2F3*), DNA replication (e.g., *PCNA*), and the p53 pathway (e.g., *MDM2*). Gene Ontology (GO) biological processes enrichment analysis for the top hits, showing the best correlation with *NUCKS1*, reveals significant enrichment for genes associated with cell cycle progression (Supplementary Fig. 1A).

In a panel of the putative cell cycle targets, *SKP2* displays the strongest and most reproducible downregulation upon NUCKS1 depletion (Supplementary Fig. 1B), and, given its role in cell cycle progression, DNA replication, and the DDR^44,46,48,49^, we focused on *SKP2*. To gain a more comprehensive understanding of *NUCKS1*’s correlation with *SKP2*, we interrogated samples from The Cancer Genome Atlas (TCGA) database. Across a range of cancer types, mRNAs encoding *NUCKS1* and *SKP2* display a significant positive correlation (Fig. 1A). There is no such correlation between *NUCKS1* and the housekeeping genes used as negative controls, *B2M* and *GAPDH* (Supplementary Fig. 1C). In particular, the correlation between *NUCKS1* and *SKP2* is most striking in glioblastoma, kidney renal papillary cell carcinoma, skin cutaneous melanoma, and uveal melanoma (Supplementary Fig. 1D).

**Fig. 1:**
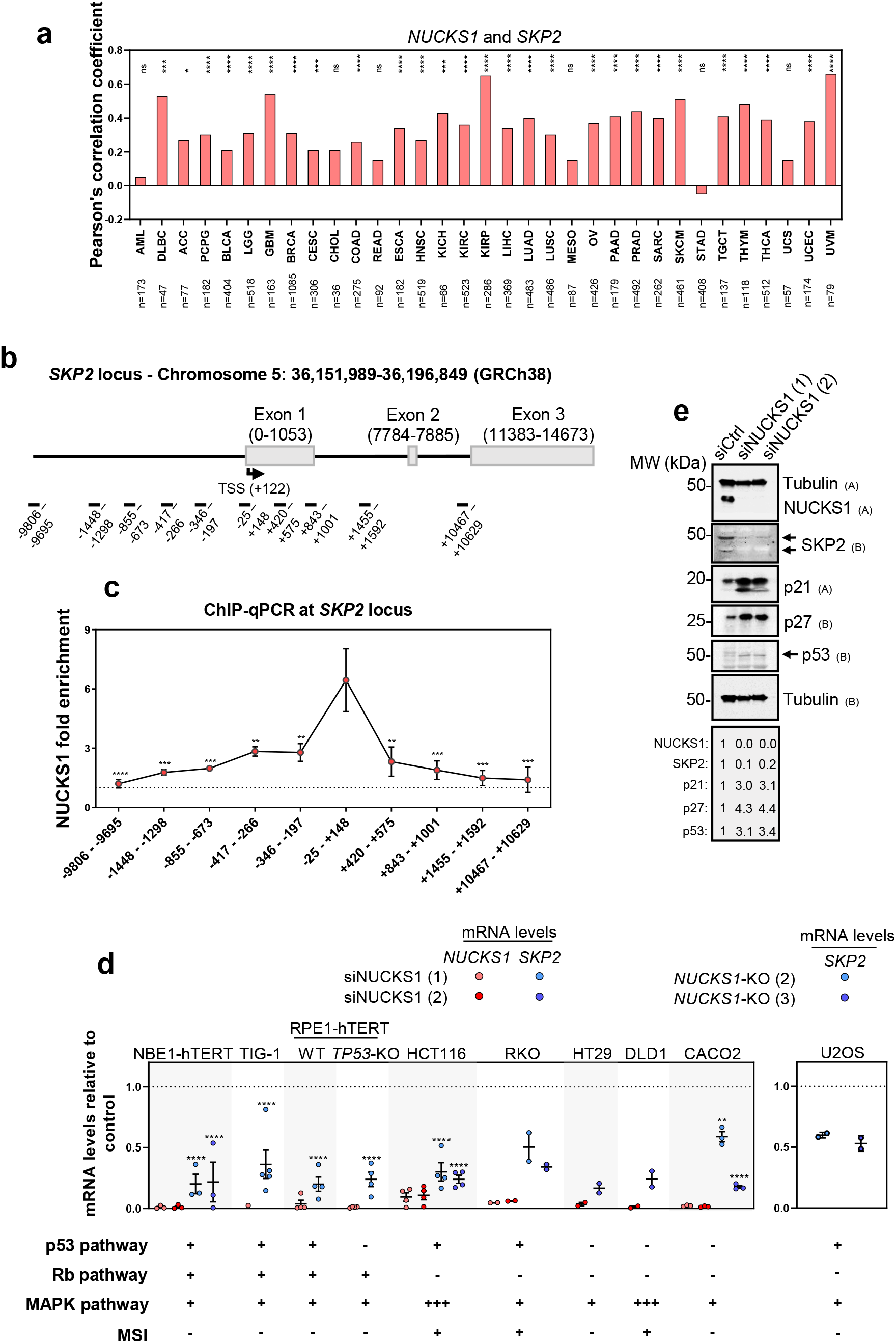
NUCKS1 transcriptionally controls the SKP2-p21/p27 axis. A) Pearson’s correlation (two-tailed) of *NUCKS1* and *SKP2* mRNAs from TCGA datasets, made using data from GEPIA2^70^. B) Map of human *SKP2* promoter annotated with sequence positions of ChIP-qPCR primers. C) ChIP-qPCR of NUCKS1 on the *SKP2* promoter in U2OS cells. Ordinary one-way ANOVA with Dunnett’s multiple comparisons test, using -25 - +148 as a reference. Data are presented as mean +/- SEM from 3 independent experiments. p-values are in order as follows: 0.0001, 0.0004, 0.0007, 0.0058, 0.0051, 0.0016, 0.0006, 0.0002, 0.0002. D) Left: RT-qPCR after control or siRNA-mediated NUCKS1 depletion. The dotted line denotes mRNA levels in siCtrl-treated cells. Right: RT-qPCR in two different clones of *NUCKS1*-KO U2OS cells. The dotted line denotes mRNA levels in WT U2OS cells. Left: One-way ANOVA with Sidak multiple comparisons test. Right: One-way ANOVA with Dunnett’s multiple comparisons test. Data are presented as mean +/- SEM from 1-4 independent experiments. p-values are in order as follows: <0.0001, <0.0001, <0.0001, <0.0001, <0.0001, <0.0001, <0.0001, 0.0015, <0.0001. E) Western blot in control- or NUCKS1-depleted RPE1-hTERT cells. Representative of 3 independent experiments. MW: molecular weight, kDa: kilodaltons. Source data are provided as a source data file.

To confirm binding of NUCKS1 at the *SKP2* promoter^16^, and to map the binding site, we designed ChIP-qPCR assays employing a panel of 10 primer sets spanning sequential regions of the *SKP2* promoter (Fig. 1B). In these assays, we found that NUCKS1 displays specificity for the chromatin directly upstream of the *SKP2* transcription start site (TSS), consistent with its role as a transcription factor (Fig. 1C).

Next, we tested the effect of NUCKS1 loss by siRNA-mediated depletion or CRISPR/Cas9-mediated deletion on *SKP2* mRNA levels (Fig. 1D, Supplementary Fig. 1E). We found that loss of NUCKS1 reduces *SKP2* gene expression across a cell line panel comprising three non-cancer cell lines (hTERT-immortalised bronchial epithelial cells: NBE1-hTERT; normal primary embryonic fibroblasts: TIG-1; hTERT-immortalised retinal epithelial cells: RPE1-hTERT), and six cancer cell lines (five colorectal cancer cell lines: HCT116, RKO, HT29, DLD1, CACO2; and osteosarcoma cells: U2OS) (Fig. 1D, Supplementary Fig. 1E). Loss of *SKP2* occurs independently of the p53 pathway, the Rb pathway, the mitogen-activated protein kinase (MAPK) pathway, and microsatellite instability (MSI) status (Fig. 1D). Furthermore, NUCKS1 depletion reduces SKP2 protein levels and increases levels of SKP2’s degradation targets, p21 and p27, confirming loss of SKP2 activity (Fig. 1E), and this is independent of p53 (Supplementary Fig. 1F, G). Consistent with the reduction in SKP2 levels, loss of NUCKS1 increases the stability of both p21 and p27, measured using cycloheximide chase assays (Supplementary Fig. 1H).

*SKP2* mRNA levels are low in early G1 and increase during the G1/S transition^50^. To test G1 cell cycle enrichment in NUCKS1-depleted cells (demonstrated in Figure 3) as an indirect mechanism for *SKP2* downregulation, we measured *SKP2* levels in cells synchronised to G0/G1 before NUCKS1 depletion (Supplementary Fig. 1I). Under these conditions, loss of NUCKS1 still reduces *SKP2* mRNA levels (Supplementary Fig. 1J), comparable with NUCKS1 depletion from asynchronous cells. These results indicate that indirect cell cycle changes do not account for reduced levels of *SKP2* in NUCKS1-depleted cells.

Altogether, these data identify *SKP2* as a transcriptional target of NUCKS1 and show that NUCKS1 regulates *SKP2* expression independently of genetic background, and in multiple cellular contexts.

### NUCKS1 levels and chromatin-binding are induced in late G1 to promote *SKP2* expression and G1/S progression

To determine whether NUCKS1 itself is subject to cell cycle-dependent regulation, and to determine the point in the cell cycle during which NUCKS1 regulates *SKP2*, we measured protein levels of NUCKS1 and SKP2 over the course of the cell cycle after release from G0/G1 synchronisation by contact inhibition. Using cyclin A2 as a marker for the onset of S phase^51^, we found that levels of NUCKS1 are low at the start of G1, increasing as cells progress into S phase (Fig. 2A). The upregulation of SKP2 (but not NUCKS1) is driven partially^52^ by an increase in its mRNA levels, which is NUCKS1-dependent (Fig. 2B). Furthermore, we detected recruitment of NUCKS1 to chromatin following release from contact inhibition-mediated G0/G1 arrest, using PCNA and MLH1 - both of which are recruited to chromatin once S phase has started^53,54^ - as controls (Supplementary Fig. 2A). The major positive stimulus for S phase entry is provided by mitogens, which activate growth factor signalling^55^. We found that stimulation of cells with mitogens following 48 h of their withdrawal triggers the recruitment of NUCKS1 to chromatin, demonstrating a potential activation of NUCKS1 by mitogenic signalling (Fig. 2C).

**Fig. 2:**
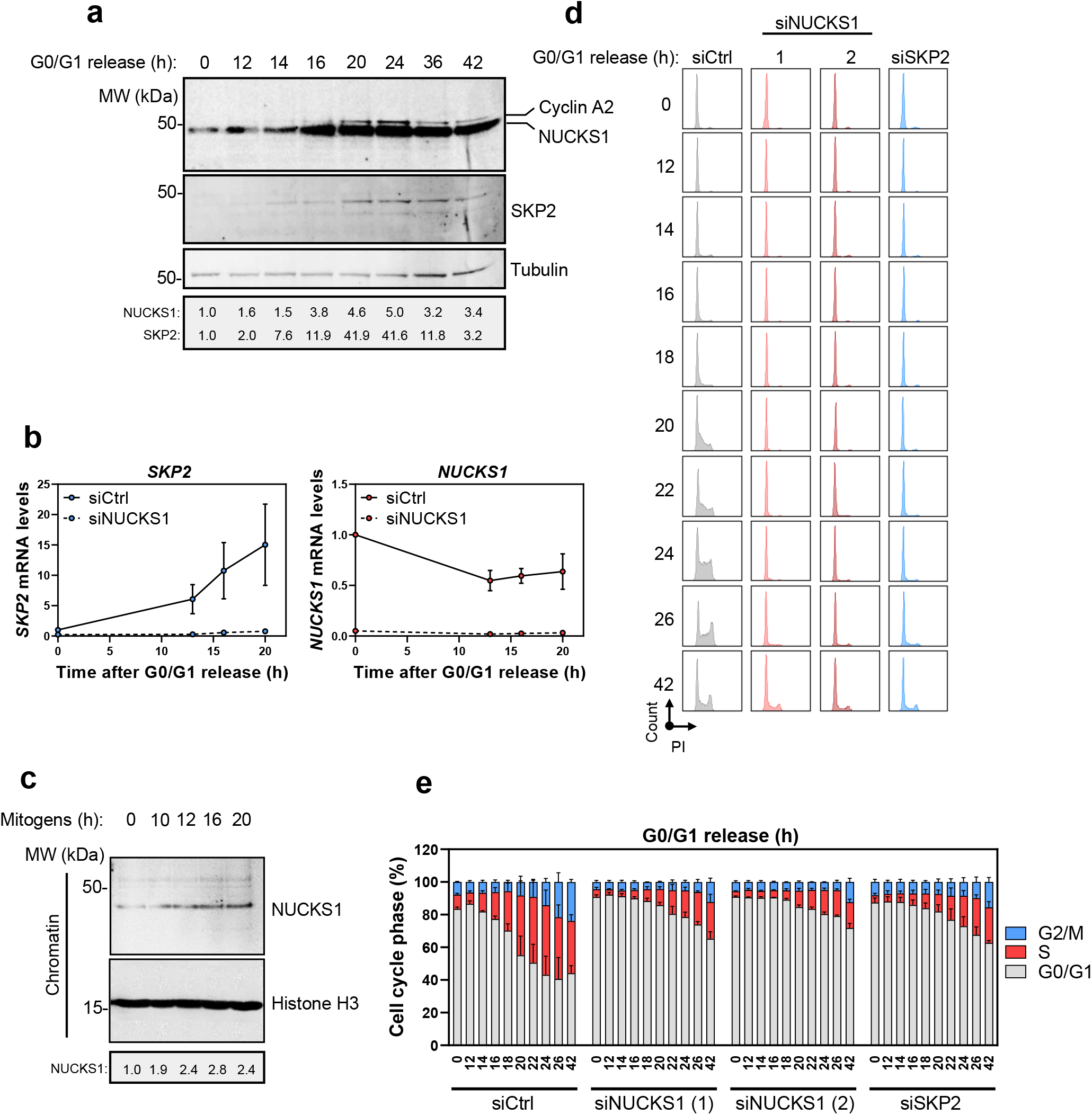
NUCKS1 levels and chromatin-binding are induced in late G1 to promote *SKP2* expression and G1/S progression. A) Western blot in whole cell extracts of RPE1-hTERT cells synchronised to G0/G1 by 72 h contact inhibition (*t*=0) followed by re-plating at low density to release cells into S phase. Representative of 3 independent experiments. B) RT-qPCR in NBE1-hTERT cells treated as in A. Data are presented as mean +/- SEM from 3 independent experiments. C) Western blot in the chromatin fraction of NBE1-hTERT cells starved of serum for 48 h (*t*=0) followed by mitogenic stimulation (15% FBS) for the indicated periods of time. Representative of 2 independent experiments. D) PI cell cycle profiles of control, NUCKS1-, or SKP2-depleted RPE1-hTERT cells treated as in A. Representative of 3 independent experiments. E) Quantification of D. Data are presented as mean +/- SEM from 3 independent experiments. MW: molecular weight, kDa: kilodaltons, PI: propidium iodide. Source data are provided as a source data file.

Stimulation of NUCKS1 during G1 progression and by mitogens suggests an active role for NUCKS1 in S phase entry. To test this, we released control, NUCKS1-, or SKP2-depleted cells from G0/G1, and measured their ability to enter S phase. We found that siRNA-mediated NUCKS1 depletion substantially delays S phase entry following G0/G1 release, phenocopying SKP2 loss (Fig. 2D, E; Supplementary Fig. 2B). Similarly, deletion of *NUCKS1* from U2OS cells impairs S phase entry (Supplementary Fig. 2C).

Together, these results demonstrate that NUCKS1’s recruitment to chromatin is stimulated by mitogens and increases during G1 progression. At the chromatin, NUCKS1 is required to induce *SKP2* transcription and S phase entry.

### NUCKS1 controls S phase entry through the SKP2-p21/p27 axis

Next, we investigated the phenotypic impact of control of the SKP2-p21/p27 axis by NUCKS1. We found that CRISPR/Cas9-mediated deletion of *NUCKS1* enriches cells in G0/G1 phase of the cell cycle, with a concomitant reduction in replicating cells (Fig. 3A). This phenotype is reversed through overexpression of wildtype NUCKS1, but not by a DNA-binding defective mutant of NUCKS1 (in which the GRP motif is mutated to AAA), confirming that NUCKS1’s DNA-binding activity is important for its role in cell cycle progression (Fig. 3B, Supplementary Fig. 3A). Furthermore, overexpression of NUCKS1 rescues cell cycle progression in NUCKS1-depleted HCT116 cells (Supplementary Fig. 3B, C), and NUCKS1 depletion delays cell cycle progression in TIG-1, NBE1-hTERT, and RPE1-hTERT cells (Supplementary Fig. 3D-F).

**Fig. 3:**
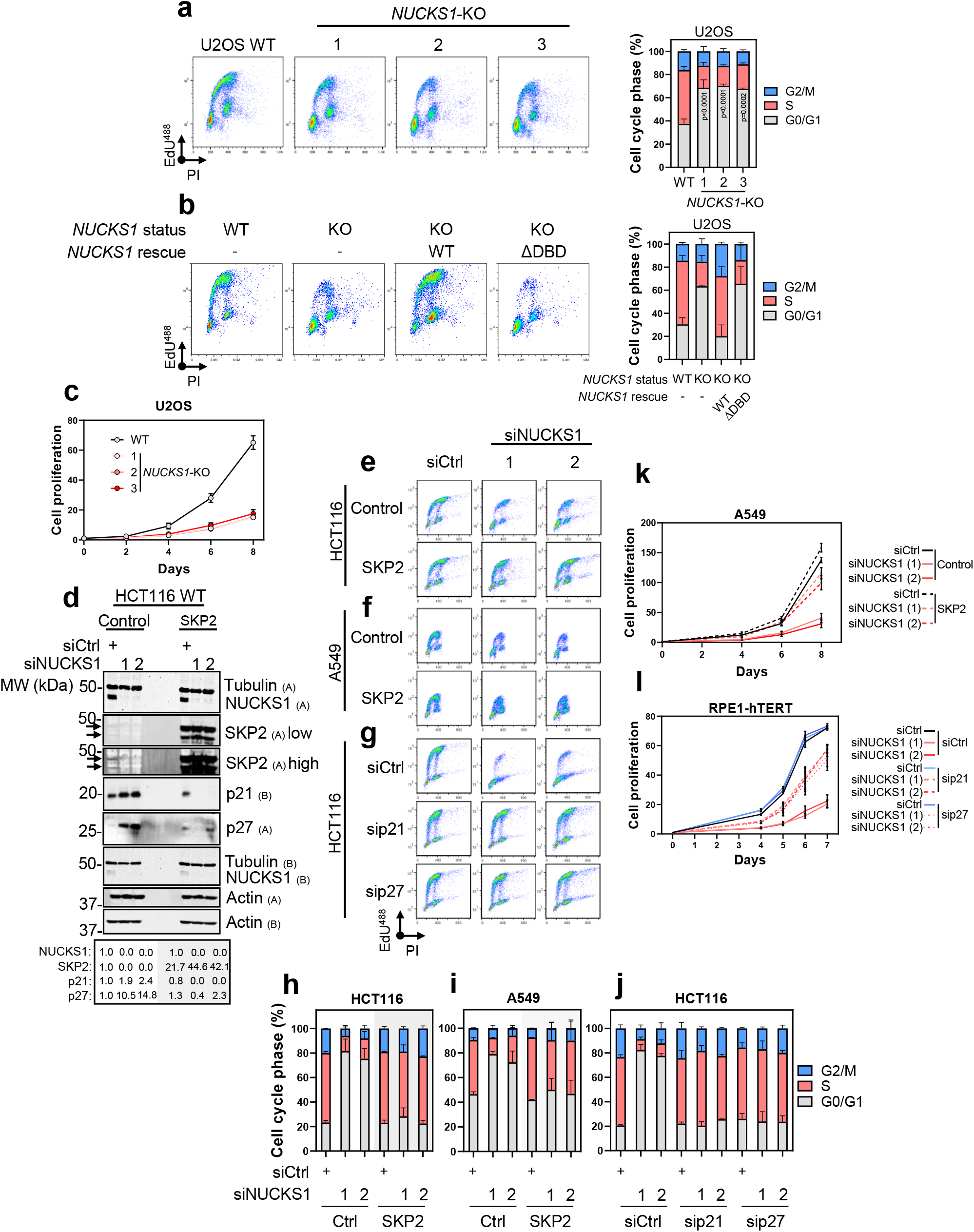
NUCKS1 controls S phase entry through the SKP2-p21/p27 axis. A) EdU/PI cell cycle profiles of WT U2OS cells and three different clones of *NUCKS1*-KO cells (left) and corresponding quantifications (right). Ordinary one-way ANOVA with Dunnett multiple comparisons test on S phase population. B) EdU/PI cell cycle profiles of WT and *NUCKS1*-KO U2OS cells expressing the indicated variants of *NUCKS1* (left) and corresponding quantifications (right). C) Proliferation assay in WT U2OS cells and three different clones of *NUCKS1*-KO cells. D) SKP2 overexpression largely rescues p21/p27 accumulation in NUCKS1-depleted HCT116 cells, measured by Western blot. E) SKP2 overexpression largely rescues HCT116 EdU/PI cell cycle profiles following treatment with control or NUCKS1 siRNA. F) SKP2 overexpression largely rescues EdU/PI cell cycle profiles in A549 cells treated with control or NUCKS1 siRNA. G) EdU/PI cell cycle profiles of HCT116 cells treated with control, p21, p27, or NUCKS1 siRNAs. H) Quantification of HCT116 SKP2 cell cycle profiles in E. I) Quantification of A549 SKP2 cell cycle profiles in F. J) Quantification of HCT116 p21/p27 cell cycle profiles in G. K) SKP2 overexpression partially rescues proliferation following NUCKS1 depletion in A549 cells. L) Proliferation assay in RPE1-hTERT cells treated with control, NUCKS1, p21 or p27 siRNAs. In A (left), B (left), D, E, F, and G, data are representative of 3 (A, D) or 2 (B, E, F, G) independent experiments. In A (right), B (right), C, H, I, J, K, and L, data are presented as mean +/- SEM from 3 (A, C, K, L) or 2 (B, H, I, J) independent experiments. MW: molecular weight, kDa: kilodaltons, PI: propidium iodide. Source data are provided as a source data file.

As a consequence, *NUCKS1* deletion from U2OS cells (Fig. 3C), and NUCKS1 depletion from TIG-1 or NBE1-hTERT cells (Supplementary Fig. 3G, H), considerably reduce cellular proliferation. NUCKS1 depletion does not cause DNA damage, measured by alkaline comet assay (which detects single-strand breaks (SSBs) and DSBs) (Supplementary Fig. 3I) or γH2AX/53BP1 immunofluorescence (Supplementary Fig. 3J), demonstrating that these phenotypes are not explained by DNA damage-induced quiescence.

We then tested whether the accumulation of p21/p27 in NUCKS1-depleted cells is due to the loss of SKP2. We found that overexpression of SKP2 in NUCKS1-depleted HCT116 (Fig. 3D) and A549 cells (Supplementary Fig. 3K) mostly induces degradation of the p21/p27 that accumulate in these cells. Consequently, overexpression of SKP2 in NUCKS1-depleted HCT116 (Fig. 3E, H) or A549 cells (Fig. 3F, I) largely rescues S phase entry. Similarly, co-depletion of SKP2’s degradation targets, p21 or p27 (Fig. 3G, J; Supplementary Fig. 3L, M, N), largely reverses cell cycle arrest.

Exploring these phenotypes further through proliferation assays, we found that SKP2 overexpression (Fig. 3K) or p21/p27 co-depletion (Fig. 3L) partially rescues the proliferation defects of NUCKS1-depleted cells. Finally, depletion of NUCKS1 from SKP2-depleted cells has no additional effect on proliferation, supporting the idea that SKP2 is a major determinant of NUCKS1’s effect on proliferation (Supplementary Fig. 3O).

These results demonstrate that NUCKS1 controls p21/p27 levels, cell cycle progression, and proliferation through its transcriptional stimulation of the *SKP2* gene, and identify the NUCKS1-SKP2-p21/p27 axis as a driving pathway for the G1/S transition.

### Analysis of NUCKS1 binding at the *SKP2* promoter

To more comprehensively understand the regulation of *SKP2* by NUCKS1, we employed the electrophoretic mobility shift assay (EMSA), using a fluorescent *SKP2* promoter probe (Fig. 4A). To do this, we started by purifying NUCKS1 from Sf9 insect cells, which preserves NUCKS1’s post-translational modifications (Fig. 4B). We found that in-tact, phosphorylated NUCKS1 displays a low affinity for the *SKP2* probe. However, dephosphorylation of NUCKS1 (using lambda phosphatase) increases the affinity of NUCKS1 for the *SKP2* probe almost 10-fold (Fig. 4C, D).

**Fig. 4:**
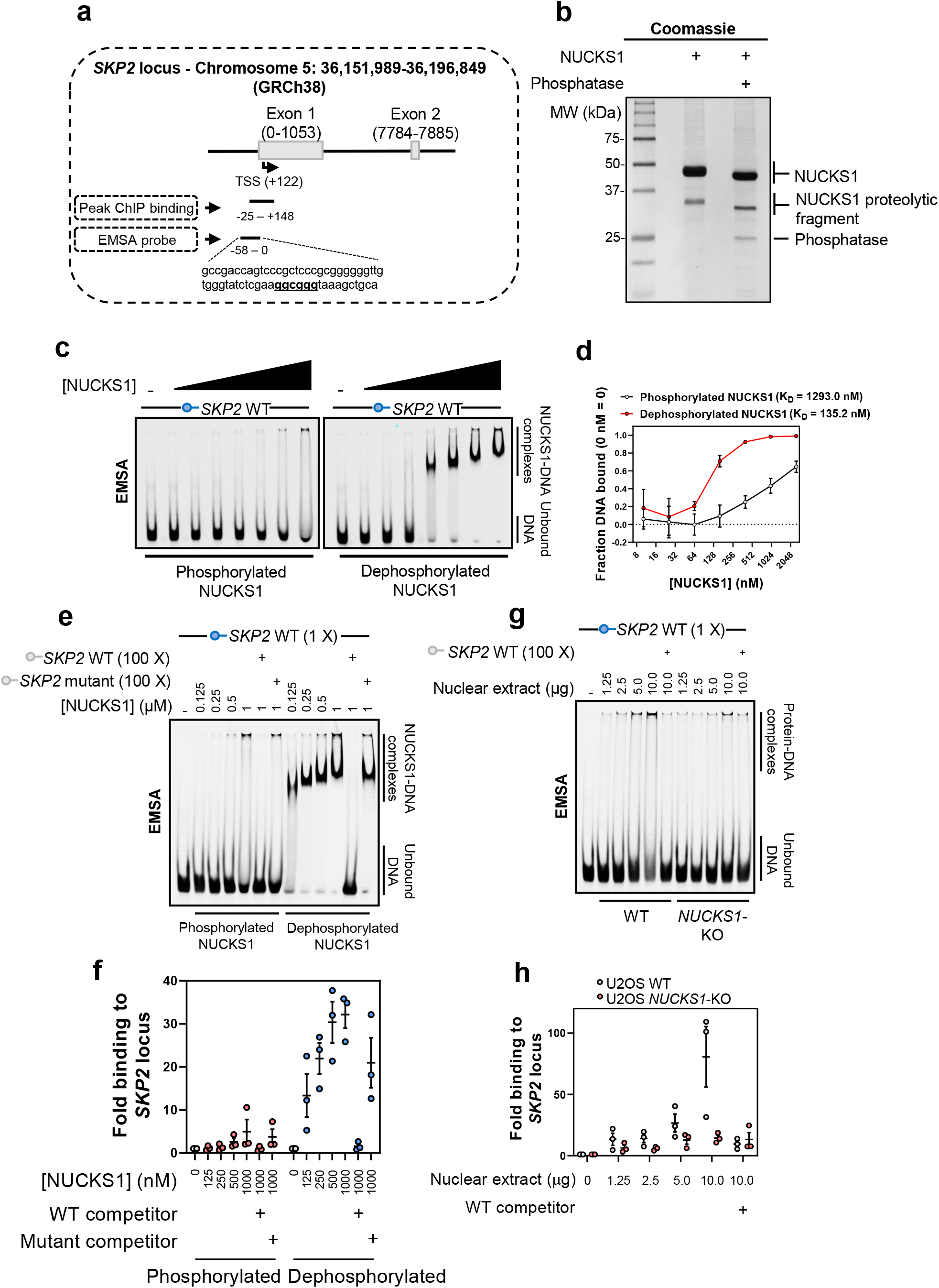
Analysis of NUCKS1 binding at the *SKP2* promoter. a) Schematic showing sequence positions of the EMSA probe in relation to *SKP2*’s transcription start site (TSS) and the region giving peak binding in our ChIP-qPCR assays. b) Coomassie gel demonstrating NUCKS1 purification. Treatment with lambda phosphatase removes NUCKS1 phosphorylation and reduces its molecular weight. c) Titration of phosphorylated or dephosphorylated NUCKS1 (10.24, 25.6, 64, 160, 400, 1000, 2500 nM) with the *SKP2* promoter probe. d) Quantification of C. e) Titration of phosphorylated or dephosphorylated NUCKS1 with the *SKP2* promoter probe. In lanes 6/12 and 7/13, respectively, 100 X molar quantity of unlabelled WT or mutant *SKP2* probe were added as competition in binding reactions. f) Quantification of E. g) Titration of WT or *NUCKS1*-KO U2OS nuclear extract with the *SKP2* promoter probe. In lanes 6 and 11, 100 X molar quantity of unlabelled WT probe was added as competition in binding reactions. h) Quantification of G. In B, C, E, and G, data are representative of 2 (B), 4 (C), or 3 (E, G) independent experiments. In D, F, and H, data are presented as mean +/- SEM from 4 (D) or 3 (F, H) independent experiments. MW: molecular weight, kDa: kilodaltons. Source data are provided as a source data file.

Since a previous publication reported a GC-box as a potential NUCKS1 binding site^16^, since there is a GC-box within the sequence of peak NUCKS1 binding to the *SKP2* promoter by ChIP-qPCR (Fig. 1C), and since recombinant NUCKS1 strongly binds the *SKP2* EMSA probe, which contains a GC-box (Fig. 4A), we mutated this sequence and performed competition EMSAs to investigate whether NUCKS1 exhibits specificity for this site. We found that the interaction of NUCKS1 with the labelled *SKP2* probe is readily outcompeted by a 100-fold excess of unlabelled WT *SKP2* probe, but not by an unlabelled mutant of the *SKP2* probe with no GC-box (Fig. 4E, F).

Finally, we performed EMSAs using WT and *NUCKS1*-KO nuclear extracts, and found that nuclear extracts from WT cells display a much higher affinity for the *SKP2* probe than extract from *NUCKS1*-KO cells (Fig. 4G, H).

Altogether, these results demonstrate that NUCKS1 directly interacts with the *SKP2* promoter’s DNA. The data suggest that this binding occurs via a GC-box in the *SKP2* promoter, and may be regulated by the phosphorylation status of NUCKS1.

### DNA damage inhibits the NUCKS1-SKP2 axis through p53-dependent transcriptional repression

DNA damage activates an ATM/p53-dependent pathway to instigate cell cycle arrest, delay DNA replication, and accomplish DNA repair^56^. To determine whether this response involves NUCKS1 or SKP2, we analysed the NUCKS1-SKP2 axis following induction of DNA damage. In U2OS cells (which express WT *TP53*, encoding p53), treatment with the chemotherapeutic drug 5-fluorouracil (5-FU) markedly reduces NUCKS1 and SKP2 protein levels, with concomitant upregulation of p21 (controlled by both p53 and SKP2), and p27 (controlled by SKP2) (Fig. 5A). 5-FU treatment also abolishes occupancy of NUCKS1 at the *SKP2* promoter (Fig. 5B), suggesting that downregulation of SKP2 is due to loss of NUCKS1 binding at its promoter. WT RPE1-hTERT cells treated with 5-FU similarly downregulate NUCKS1 and SKP2, and upregulate p21 and p27. However, this response is absent in *TP53*-KO RPE1-hTERT cells, suggesting a role for p53 in DNA damage-mediated NUCKS1/SKP2 downregulation (Fig. 5C). Consistent with loss of SKP2, the stability of p21 and p27 is extended in 5-FU-treated cells, revealed through chase assays with the translation inhibitor cycloheximide (Supplementary Fig. 4A).

**Fig. 5:**
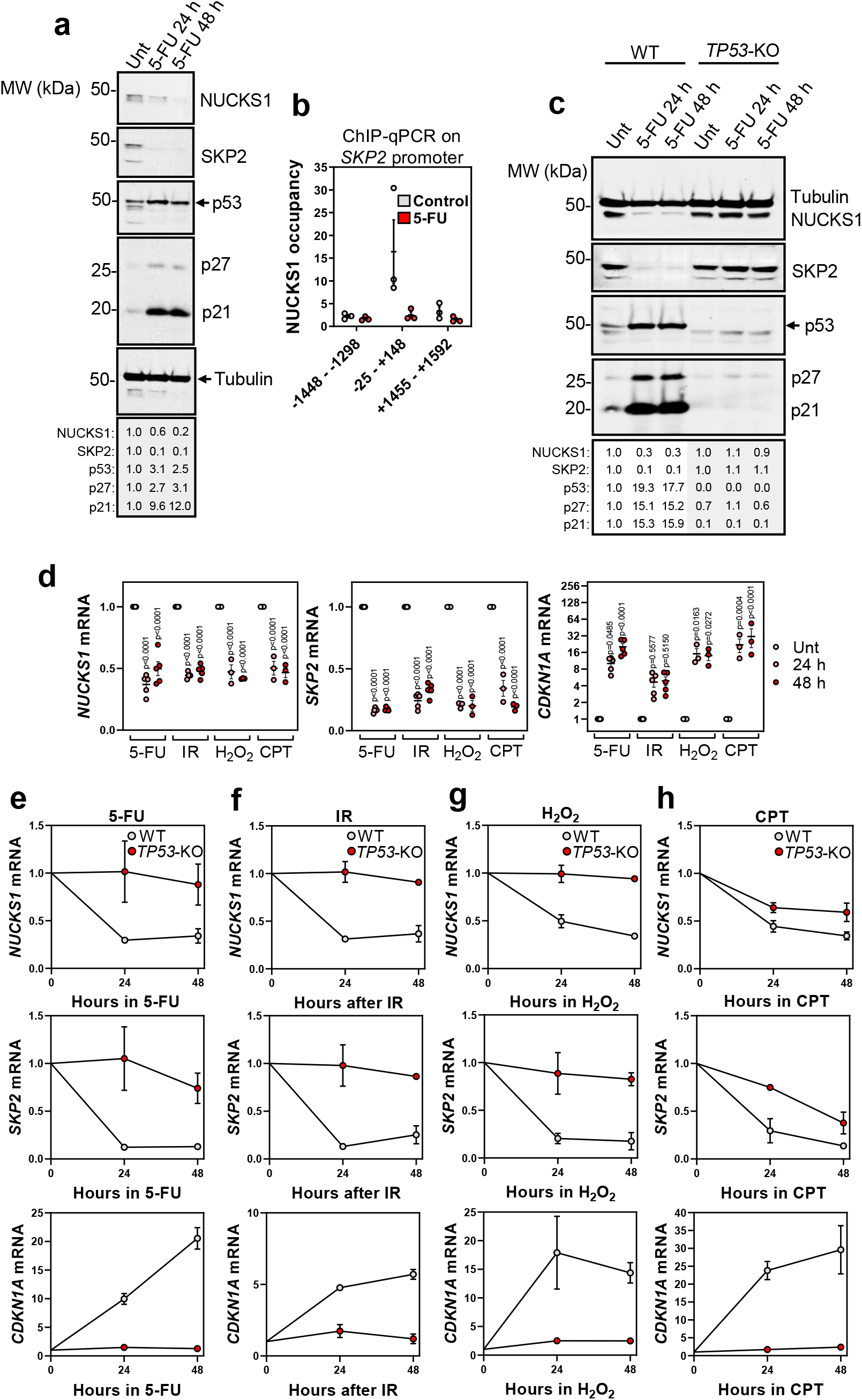
DNA damage inhibits the NUCKS1-SKP2 axis through p53-mediated transcriptional repression. a) Western blot in WT U2OS cells treated with 50 μM 5-FU for 24 or 48 h. b) ChIP-qPCR of NUCKS1 on the *SKP2* promoter in U2OS cells after 50 μM 5-FU for 48 h. c) Western blot in RPE1-hTERT WT and *TP53*-KO cells treated with 10 μM 5-FU for 24 or 48 h. d) RT-qPCR in WT RPE1-hTERT cells treated with 5-FU (10 μM), IR (4 Gy), H_2_O_2_ (200 μM), or CPT (100 nM) for 24 or 48 h. Ordinary two-way ANOVA with Dunnett’s multiple comparisons test. e) RT-qPCR after 5-FU (10 μM) in RPE1-hTERT WT or *TP53*-KO cells. f) RT-qPCR after IR (4 Gy) in RPE1-hTERT WT or *TP53*-KO cells. g) RT-qPCR after H_2_O_2_ (200 μM) in RPE1-hTERT WT or *TP53*-KO cells. h) RT-qPCR after CPT (100 nM) in RPE1-hTERT WT or *TP53*-KO cells. In A and C, data are representative of 3 independent experiments. In B, D, E, F, G, and H, data are presented as mean +/- SEM from 3 (B, H), 3-5 (D) or 2 (E-G) independent experiments. MW: molecular weight, kDa: kilodaltons. Source data are provided as a source data file.

To investigate this putative role for p53, we used RT-qPCR to measure the mRNA levels of *NUCKS1* and *SKP2* after treatment with 5-FU, IR, hydrogen peroxide (H^2^O^2^), and camptothecin (CPT), DNA-damaging agents which induce distinct DNA lesions. Upregulation of *CDKN1A* mRNA, encoding p21, was used as a control for p53 activation. We found that *NUCKS1* and *SKP2* transcripts are substantially reduced in response to all tested DNA-damaging agents (Fig. 5D). Consistent with Western blotting data, this is largely dependent on p53 (Fig. 5E-H). Induction of DNA damage also induces cell cycle changes which similarly depend on *TP53* status (Supplementary Fig. 4B). These results demonstrate that DNA damage induces a p53 response, involving downregulation of NUCKS1 and SKP2, upregulation of p21 and p27, and cell cycle arrest.

Next, we sought to understand the mechanism underpinning p53-dependent downregulation of NUCKS1 and SKP2. Much p53-mediated transcriptional repression relies on activation of the DREAM transcriptional repression complex by p53-induced p21^57^. To investigate whether this pathway is involved in the downregulation of NUCKS1 or SKP2, we treated WT and *CDKN1A*-knockout cells with 5-FU. As expected, transcripts of the p21-DREAM target *CCNB1*^58^, used as a positive control, are only reliably downregulated upon DNA damage in WT cells. However, transcripts of *NUCKS1* and *SKP2* are downregulated both in WT and *CDKN1A*-knockout cells, suggesting that *NUCKS1* and *SKP2* are not targets of the p21-DREAM pathway (Supplementary Fig. 4C-E).

Finally, we found that RNAi-mediated p53 depletion in TIG-1 cells, which express high endogenous p53 levels^59^ (Supplementary Fig. 4F), as well as deletion of *TP53* from RPE1-hTERT cells (Supplementary Fig. 4G), increases *NUCKS1* levels, suggesting that p53 may regulate *NUCKS1*/*SKP2* expression both under basal conditions, as well as following p53 activation.

We propose that the p53-NUCKS1-SKP2-p21/p27 axis constitutes a checkpoint pathway for the G1/S transition, which may respond to DNA damage to prevent the replication of damaged DNA.

### Copy number gain and p53 loss contribute to *NUCKS1* and *SKP2* overexpression in cancer

Transcriptional overexpression of *NUCKS1* and *SKP2* has been reported in numerous cancer types^18,33–36,38,46,60–62^. Although some reports, focused on specific cancer types, attribute this to increased copy number^33,37,39,46,61,63^, no pan-cancer analyses have been performed, and the full mechanisms underlying the upregulation remain poorly defined. Seeking to explore this further, we analysed *NUCKS1* and *SKP2* expression in TCGA datasets. *NUCKS1* and *SKP2* are overexpressed in most TCGA datasets, including many shared cancer types (Fig. 6A, B). Consistent with oncogenic functions for *NUCKS1*^39^ and *SKP2*^62^, both genes are subjected to copy number increase in many cancers, while deletions are rare (Fig. 6C, D). These results confirm that increased copy number of *NUCKS1* and *SKP2* can contribute to their overexpression in cancer.

**Fig. 6:**
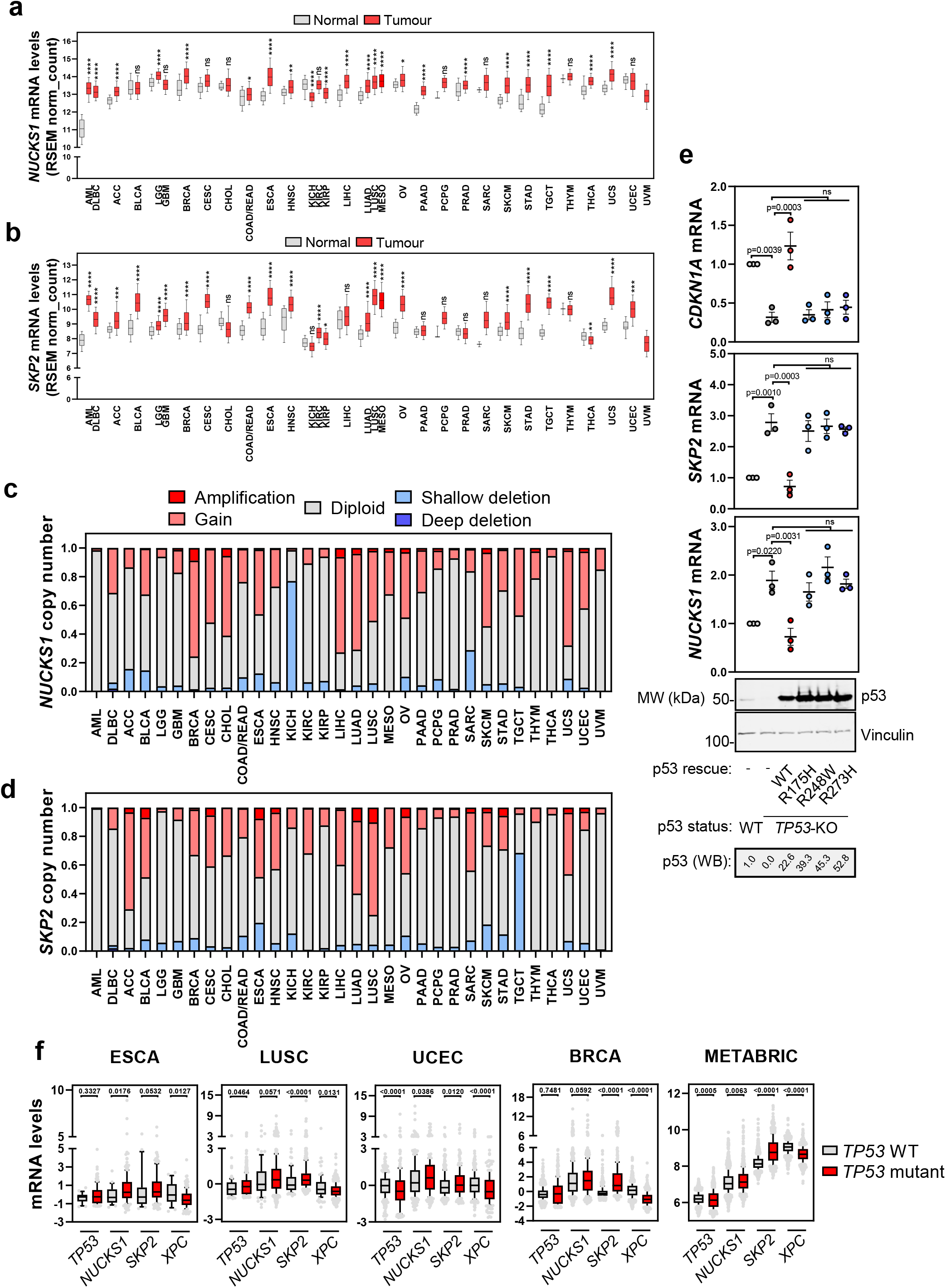
Copy number gain and p53 loss contribute to *NUCKS1* and *SKP2* overexpression in cancer. A) *NUCKS1* expression in normal vs. tumour tissue, using data from UCSC Xena^71^. In A and B, box plots show median values along with 25/75% (box) and 10/90% (whiskers). Statistics were analysed using Kruskal Wallis with Dunn’s post test. p-values are in order as follows: <0.0001, <0.0001, <0.0001, >0.9999, <0.0001, 0.1198, <0.0001, >0.9999, >0.9999, 0.0148, <0.0001, 0.0019, <0.0001, >0.9999, <0.0001, <0.0001, <0.0001, <0.0001, <0.0001, 0.0187, <0.0001, 0.6879, <0.0001, >0.9999, <0.0001, <0.0001, <0.0001, >0.9999, <0.0001, <0.0001, >0.9999. B) *SKP2* expression in normal vs. tumour tissue, using data from UCSC Xena^71^. p-values are in order as follows: <0.0001, <0.0001, 0.0002, <0.0001, <0.0001, <0.0001, <0.0001, <0.0001, >0.9999, <0.0001, <0.0001, <0.0001, >0.9999, <0.0001, 0.0476, 0.3634, <0.0001, <0.0001, <0.0001, <0.0001, >0.9999, >0.9999, >0.9999, 0.3562, <0.0001, <0.0001, <0.0001, >0.9999, 0.0023, <0.0001, 0.0006. C) *NUCKS1* copy number changes in cancer, using data from TCGA PanCancer Atlas datasets in CBioPortal^72,73^. D) *SKP2* copy number changes in cancer, using data from TCGA PanCancer Atlas datasets in CBioPortal^72,73^. E) Western blot and RT-qPCR in p53 proficient vs. deficient HCT116 cells expressing indicated variants of p53. Ordinary one-way ANOVA with Tukey’s multiple comparisons test. Upper: data are presented as mean +/- SEM from 3 independent experiments. Lower: data are representative of 3 independent experiments. ns p-values are in order as follows – *NUCKS1*: 0.8983, 0.8478, 0.9995; *SKP2*: 0.9412, 0.9984, 0.9786; *CDKN1A*: 0.9999, 0.9802, 0.9334. F) Analysis of *TP53, NUCKS1, SKP2* or *XPC* (used as a positive control for p53 activity) mRNA levels in WT vs. *TP53* mutant tumours, using PanCancer Atlas or METABRIC datasets in CBioPortal^72,73^. Units: log RNA Seq V2 RSEM (ESCA-BRCA) and mRNA expression microarray (METABRIC). Box plots show median values along with 25/75% (box) and 10/90% (whiskers) and outliers. Two-tailed Mann-Whitney test. ESCA p53 WT n=24 patients, p53 mutant n=157 patients. LUSC p53 WT n=79 patients, p53 mutant n=402 patients. UCEC p53 WT n=323 patients, p53 mutant n=192 patients. BRCA p53 WT n=717 patients, p53 mutant n=347 patients. METABRIC p53 WT 1245 patents, p53 mutant n=659 patients. MW: molecular weight, kDa: kilodaltons. Source data are provided as a source data file.

Because we found that p53 negatively regulates levels of *NUCKS1* and *SKP2* (Fig. 5), we investigated the effect of p53 mutation in cancer. To do so, we used p53-proficient vs. -deficient HCT116 cells, and expressed WT p53 as well as its DNA-binding mutants, R175H, R248W, and R273H, which frequently drive cancer^64^. We found that p53-deficient HCT116 cells have increased levels of both *NUCKS1* and *SKP2*. Notably, overexpression of WT p53 - but not its DNA-binding mutants - substantially reduces *NUCKS1*/*SKP2* levels (Fig. 6E). These results further support our finding that p53 negatively regulates levels of *NUCKS1* and *SKP2*, and demonstrate that p53 mutants defective for DNA-binding lose the ability to repress *NUCKS1*/*SKP2*.

Finally, we asked whether p53 mutations also affect *NUCKS1*/*SKP2* expression in cancer patients, using TCGA datasets. Consistent with our *in vitro* data, we found that mutation of p53 correlates with overexpression of *NUCKS1*/*SKP2* in several cancer types (Fig. 6F).

Together, these results show that increased copy number, as well as p53 mutation, contribute to the overexpression of *NUCKS1* and *SKP2* in many cancers. This may enable cancer cells to proliferate in the absence of mitogenic stimulation, or in the presence of DNA damage.

## Discussion

Here, we identify the NUCKS1-SKP2-p21/p27 axis as a cell cycle checkpoint pathway, which responds antagonistically to mitogen and DNA damage input to control S phase entry. In early G1 cells and in the absence of mitogens, NUCKS1 protein levels and chromatin retention are low, ensuring its inhibition in non-replicating cells. NUCKS1 is upregulated and recruited to chromatin during G1/S progression, permitting NUCKS1 to stimulate the expression of *SKP2*, the F-box component of the SCF^SKP2^ ubiquitin ligase, leading to the degradation of p21/p27 and S phase entry. In contrast, DNA damage induces p53-dependent transcriptional repression of *NUCKS1*, leading to loss of SKP2 and upregulation of p21/p27 for cell cycle arrest. Some cancer cells hijack this mechanism, increasing *NUCKS1/SKP2* copy numbers and transcriptionally upregulating *NUCKS1* and *SKP2* through p53 mutation. We propose that this may enable cancer cells to sustain proliferation, even in the absence of mitogens or in the presence of DNA damage (Fig. 7).

**Fig. 7:**
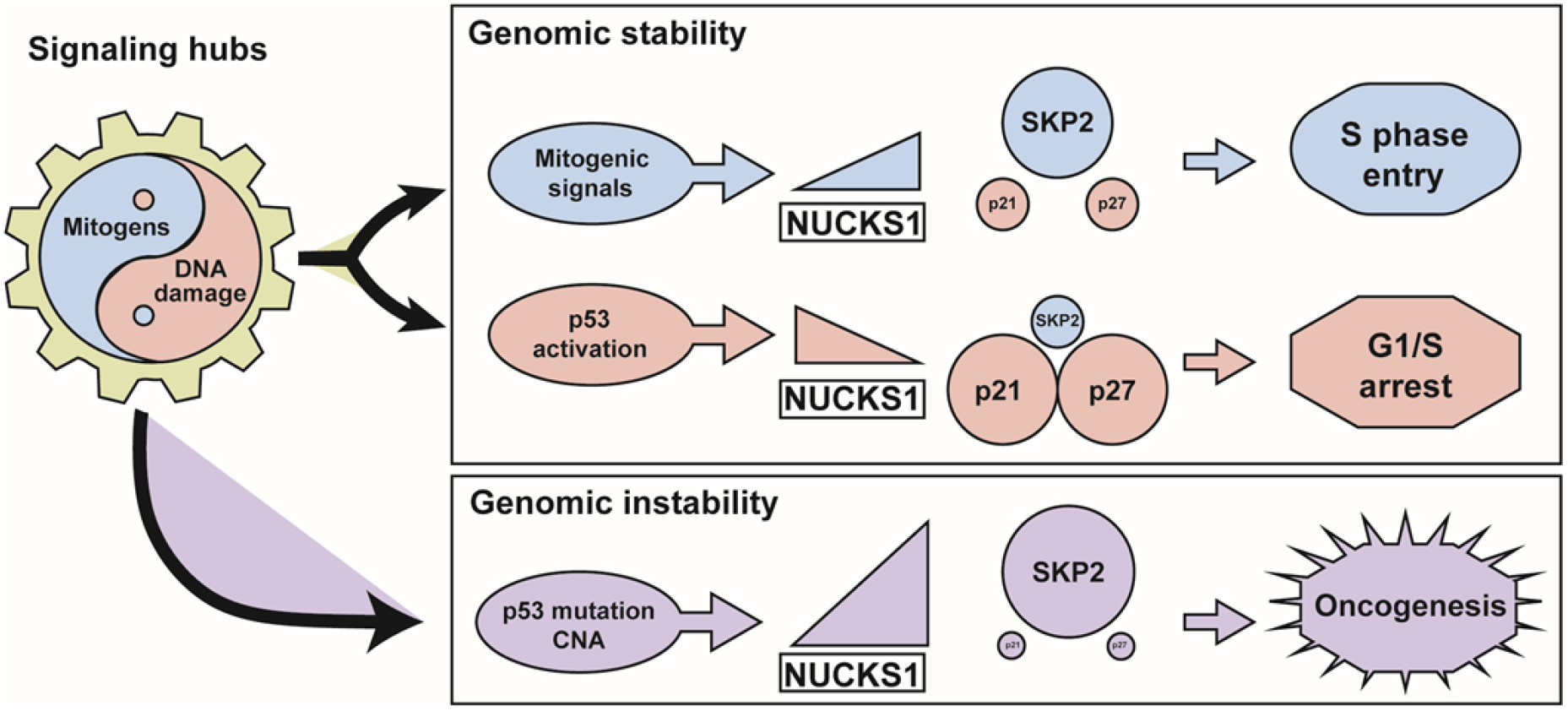
Model depicting the role for NUCKS1 in the S phase entry decision. The NUCKS1-SKP2-p21/p27 axis constitutes a signalling hub which integrates the opposing cell cycle signals, mitogens and DNA damage. Mitogens stimulate binding of NUCKS1 to the *SKP2* promoter, *SKP2* expression, p21/p27 degradation, and S phase entry. DNA damage induces p53-dependent repression of *NUCKS1*, leading to *SKP2*’s transcriptional downregulation, upregulation of p21/p27, and cell cycle arrest. Some cancer cells increase *NUCKS1/SKP2* copy number and mutate p53, leading to *NUCKS1* and *SKP2* overexpression. CNA: copy number alteration.

Our study identifies the SKP2-p21/p27 pathway as the second pathway transcriptionally controlled by NUCKS1, after the insulin receptor pathway^16^. However, the question of precisely how NUCKS1 regulates transcription remains unanswered. NUCKS1 is known to bind chromatin with higher affinity than naked DNA, does not bind ssDNA, and binds D-loops better than dsDNA^19^. NUCKS1 does not have a transcription activation domain, but promotes chromatin accessibility at - and recruits RNAPII to – its target promoters^16^. It is possible that NUCKS1 cooperates with other transcription factors to direct transcription; for example, NUCKS1 has been reported as an activator of NF-κB^65^. Since NF-κB regulates *SKP2* levels^66^, NUCKS1 may cooperate with NF-κB to control *SKP2* expression. Nevertheless, future work will focus on characterising NUCKS1’s interactome, to more deeply investigate its mechanism for transcriptional regulation.

Our EMSA data (Fig. 4) and others’ ChIP-Seq data^16^ reveal that NUCKS1 displays affinity for the GC-box target sequence. The sequence we identify, GGCGGG, is present twice within the 600 nucleotide *SKP2* promoter, but is absent from the remaining ∼45,000 nucleotides of the *SKP2* gene, which may explain the specificity of NUCKS1 for the *SKP2* promoter *in vivo*, and for other NUCKS1 targets more broadly. Going forward, research should focus on the structural basis of NUCKS1’s interaction with the GC-box, and investigate whether NUCKS1 has multiple target DNA-binding sequences.

We show that p53 mediates the transcriptional downregulation of *NUCKS1* in response to DNA damage (Figure 5), but we do not fully characterise the mechanism. Binding of p53 at the *NUCKS1* promoter, with enrichment following DNA damage, has been detected as part of genome-wide ChIP-Seq studies^32^, and our data showing that the downregulation of *NUCKS1* following DNA damage is independent of p21-DREAM (Supplementary Fig. 4) suggest that *NUCKS1*’s repression may be a direct result of p53 binding. To explore this further, it would be useful to measure the rate of synthesis of new *NUCKS1* transcripts, as well as the stability of *NUCKS1* transcripts, to determine whether p53 controls *NUCKS1*’s transcription itself or the stability of its mRNA. Complementary luciferase assays using the *NUCKS1* promoter could also reveal whether p53 controls the activity of the *NUCKS1* promoter. Furthermore, would mutation of a putative binding site for p53 in the *NUCKS1* promoter alter *NUCKS1* expression, DNA damage resistance, cell cycle progression, and proliferation? These experiments will form part of future studies.

NUCKS1 is the most post-translationally modified protein in the human proteome (for its size) and its major modification is phosphorylation^23^. NUCKS1 is phosphorylated by the G2/M cell cycle kinase CDK1 at S181, reducing NUCKS1 binding to DNA^26,27^, although the *in vivo* function of this phosphorylation is not completely understood. Interestingly, S181 phosphorylation of NUCKS1 could act to reset the level of chromatin-bound NUCKS1 for the daughter G1 phase, during which CDK1 activity is low, and explain the G1/S chromatin recruitment of NUCKS1 that we observe (Supplementary Fig. 2). Furthermore, the DDR kinase ATM promotes the indirect phosphorylation of NUCKS1 at S181 following DNA damage^19,29,67^. Therefore, ATM-dependent NUCKS1 phosphorylation could provide a secondary mechanism to p53-dependent transcriptional repression, to ensure NUCKS1’s removal from cell cycle promoters after DNA damage, and warrants investigation in the future. Notably, phosphorylation of NUCKS1 at S181 may also explain our EMSA data, which reveal a significant increase in DNA-binding affinity following NUCKS1 dephosphorylation (Figure 4).

By stimulating the activity of RAD54, NUCKS1 promotes HR, the S/G2-specific DSB repair pathway^19,22^, demonstrating that NUCKS1 acts to maintain the fidelity of DNA replication. Consistent with this, we show that NUCKS1 levels remain high throughout S phase and into G2 (Fig. 2). These findings raise a model in which NUCKS1 stimulates entry into S phase and promotes the fidelity of the ensuing DNA replication through HR, after its role in S phase entry is achieved. Notably, this function would mirror that of other G1/S factors, which boost both S phase entry and DNA repair, including E2F1^68^ and SKP2 itself^49^.

In summary, our study identifies NUCKS1 as an important factor for the G1/S transition, placing NUCKS1 within the SKP2-p21/p27 axis. Future studies will investigate NUCKS1’s mechanism of transcriptional regulation, mechanisms for its regulation by posttranslational modification, and delve deeper into its roles in oncogenesis.

## Author contributions

G.L.D. and A.J.L. conceived the study and were in charge of overall direction and planning. S.H., C.P.G., P.L. and A.J.L. performed experiments. All authors designed and analysed experiments. V.D. gave critical suggestions and provided reagents. A.J.L., K.R. and G.L.D. supervised the project. S.H., A.J.L., K.R. and G.L.D. wrote the manuscript. All authors read and approved the manuscript.

## Acknowledgements

The authors thank Prof. I. Cheeseman (Department of Biology, Massachusetts Institute of Technology), Dr. R. Chapman (Nuffield Department of Medicine, University of Oxford) and Prof. S. Maheswaran (Massachusetts General Hospital Cancer Center, Harvard Medical School) for providing cell lines. We thank Prof. E. O’Neill (Department of Oncology, University of Oxford) for providing plasmids, Prof. A. Østvold (Department of Biochemistry, University of Oslo) for NUCKS1 antibodies, and Dr. S. Mukhopadhyay, Prof. N. Burgess-Brown (Structural Genomics Consortium, University of Oxford), and Dr. S. Khoronenkova (Department of Biochemistry, University of Cambridge) for their help with NUCKS1 purification. The authors thank present and past members of the Dianov and Ramadan labs for discussions and technical help.

G.L.D. is supported by grants from the Medical Research Council [H3RWGJ00.H302.1], Cancer Research UK [C5255/A15935], and the Russian Science Foundation grant (№19-74-20069). K.R. is supported by the Medical Research Council Programme (MC_PC-12001/1 and MC_UU-00001/1) and Breast Cancer Now (2019DecPR1406). S.H. was supported by the Radcliffe-Oncology Studentship at University College, University of Oxford.

## Materials and Correspondence

Correspondence and material requests should be addressed to K.R.

## Data availability

Source data are provided with this paper.

Supplementary Fig. 1A uses the SEEK database^47^ (http://seek.princeton.edu/) and Genecodis3^69^ (http://genecodis.cnb.csic.es). Fig. 1A and Supplementary Fig. 1C and 1D were generated using data from GEPIA2^70^ (http://gepia2.cancer-pku.cn/#index). Fig. 6A and B were generated using data from UCSC XENA (https://xena.ucsc.edu/), using RSEM norm count values from GTEX/TCGA normal datasets, and TCGA tumour datasets^71^. Fig. 6C, D and F were generated using data from CBioPortal^72,73^ (https://www.cbioportal.org/).

## Methods

### Cell culture

Cell lines (Supplementary Table 1) were cultured in DMEM (Life Technologies) with 15% (TIG-1, NBE1-hTERT^74^) or 10% (U2OS, U2OS *NUCKS1*-KO, RPE1-hTERT, RPE1-hTERT *TP53*-KO, RPE1-hTERT *CDKN1A*-KO^75^, HT29, A549 SKP2 doxycycline-inducible^76^, DLD1, RKO, HCT116, HCT116 *TP53*-KO^77^, CACO2) FBS, at 37°C in a humidified atmosphere with 5% CO_2_. All cells tested negative for mycoplasma. For ionising radiation, treatments were performed using a GSR-D1 137Cs γ-irradiator (RPS Services) at a dose rate of 1.8 Gy/min.

### siRNA and plasmid transfections

siRNA transfections were performed using Lipofectamine RNAiMAX, according to the manufacturer’s instructions. Cells were transfected with 30 nM siRNA for 3-7 days. siRNA sequences used are as follows:

siCtrl: Eurogentec, SR-CL000-005; siNUCKS1 (1): GAGGCGAUCUGGAAAGAAU; siNUCKS1 (2): GGCAUCUAAAGCAGCUUCU; siNUCKS1 (3’ UTR): GCAGGAGGGACUAGAGAAAUU; siSKP2: GCUUCACGUGGGGAUGGGA; sip21: GAUGGAACUUCGACUUUGU; sip27: AAGGUUGCAUACUGAGCCAAG; sip53: AAGACUCCAGUGGUAAUCUAC.

Plasmid transfections were performed using Lipofectamine 3000, according to the manufacturer’s instructions. Assays were performed 48 h after plasmid transfection. Plasmids used in the study are listed in Supplementary Table 2.

### CRISPR-Cas9 genome editing

*NUCKS1* CRISPR/Cas9 KO plasmid (sc-413018) and *NUCKS1* HDR plasmid (sc-413018-HDR) were co-transfected into early passage U2OS cells. Cells were treated with 5 μg/ml puromycin for 3 days to select successfully-transfected cells, and seeded as single cells. Colonies were expanded and successful clones were confirmed using RT-qPCR and Western blotting.

### Western blotting

Whole cell extracts were prepared as described previously^78^. Nuclear/chromatin fractionations were performed as described previously^79^. Proteins were resolved using SDS-PAGE and transferred onto Immobilon-FL PVDF membranes (Millipore). Membranes were blocked using Odyssey blocking buffer (Li-Cor) and blotted using the antibodies indicated in Supplementary Table 3. Western blot detection was performed using the Odyssey image analysis system (Li-Cor Biosciences). Analysis and quantification were performed using Image Studio Lite Ver 5.2.

### RT-qPCR

Total RNA was extracted using the RNeasy kit (QIAGEN). Reverse transcription was performed using the SuperScript II Reverse Transcriptase kit (Thermofisher). RT-qPCR was performed using Fast SYBR Green Master Mix (Thermofisher) and the 7500 Fast Real-Time PCR System (Applied Biosystems), with the comparative CT method for quantification. Analysis was performed using 7500 Software v2.0.6. Reference genes used for RT-qPCR are *B2M*/*GAPDH*/*TBP*. Primer sequences are listed in Supplementary Table 4.

### Protein expression and purification

Production of baculoviral particles, infection of Sf9 cells, and expression of recombinant protein was performed as described previously^80^. Mid log phase *Spodoptera frugiperda* (Sf9) cells (2×10^6^/ml) were transfected with pDEST53-*NUCKS1* bacmid using Cellfectin II transfection reagent in a 6-well plate format, according to the manufacturer’s protocol. Following incubation for 5 days at 27 °C, medium containing P0 baculovirus was collected and stored at 4 °C, protected from light. Two sequential rounds of virus amplification were performed to generate higher titer P2 baculovirus stocks. Sf9 cells were infected with P2 virus (120 μg/50 ml Sf9), and incubated at 27 °C for 3 days on an orbital shaker. Cells were harvested by centrifugation (900 g, 20 min, 4 °C), washed with PBS, pelleted again, and then stored at -80 °C. Cell pellets were resuspended in buffer A (50 mM HEPES pH 7.5, 300 mM NaCl, 5% (w/v) glycerol) supplemented with 1 mM TCEP, and 1:500 (v/v) protease inhibitor cocktail (Sigma-Aldrich, P8849) (12 ml buffer A per 100-ml culture cell pellet), and lysed by sonication, followed by incubation with Benzonase (20 U/μl) for 30 min on ice. Cell lysate were clarified by centrifugation, and supernatant passed through a 0.45-μm syringe filter. The supernatant was then supplemented with 5 mM imidazole prior to loading onto a 1-ml HisTrap column (GE Healthcare) attached to an AKTA system at 1 ml/min. After sample loading, the column was washed with buffer A containing 5 mM (10 column volumes (CV)) and 50 mM imidazole (10CV). NUCKS1 was eluted with a linear 50-250 mM imidazole gradient (20CV) and 0.5 ml fractions were collected. His_6_-tagged NUCKS1-containing fractions were pooled, and dialysed against storage buffer (50 mM Tris pH 7.5, 150 mM NaCl, 5% (w/v) glycerol, 0.5 mM DTT). To improve purity, His_6_-tagged NUCKS1 was further purified by size-exclusion chromatography. 1.5 mg of HisTrap-purified NUCKS1 was diluted to 200 μl in storage buffer and loaded onto a Superdex 200 HR 10/30 column (GE Healthcare, Little Chalfont, UK), and 0.5 ml fractions collected.

### Immunofluorescence

Cells seeded on coverslips were subjected to pre-extraction in a buffer containing 10 mM HEPES, 100 mM NaCl, 0.3 M sucrose, 3 mM MgCl_2_, and 0.1% Triton X-100 for two minutes, washed twice using a buffer containing 10 mM HEPES, 100 mM NaCl, M sucrose, and 3 mM MgCl_2_, and then fixed in 4% formaldehyde for 15 minutes on ice. Cells were blocked overnight in 5% BSA at 4°C, and primary antibodies (indicated in Supplementary Table 3) were diluted in 2.5% BSA and incubations performed for 1 h at RT. Cells were washed in PBS, incubated in secondary antibodies (indicated in Supplementary Table 3) for 1 h at RT, and stained with DAPI. Microscopy was performed using the Nikon NiE and quantification was performed using CellProfiler^81^.

### Site-directed mutagenesis

NUCKS1’s GRP DNA-binding motif was mutated to AAA using the Phusion Site-Directed Mutagenesis Kit (Thermofisher), with primers as follows: gcctttgaagctgtggcggcagccactttccctttgcc/ggcaaagggaaagtggctgccgccacagcttcaaaggc. Mutant *NUCKS1* was validated by sequencing.

### Comet assays

Alkaline comet assays were performed as described previously^82^, using the Nikon NiE and Andor Komet7.1 software.

### Proliferation assays

Cells were seeded at day 0, treated as indicated, and viable cells were counted at indicated days, using Trypan Blue staining (Life Technologies) and the Countess™ Automated Cell Counter (Thermo Fisher Scientific).

### EMSAs

For NUCKS1 EMSAs, recombinant NUCKS1 was dephosphorylated using lambda phosphatase (100 units/1 μg of recombinant NUCKS1), in the presence of Protein MetalloPhosphatases buffer and MnCl_2_ (1 mM), for 90 min at 30 °C, followed by addition of phosphatase inhibitor cocktail (50 x, Merck Millipore). Consequently, binding reactions using indicated quantities of intact or dephosphorylated NUCKS1, or WT or *NUCKS1*-KO nuclear extract, were set up in the presence of binding buffer (20 mM Tris-HCl pH 7.5, 100 mM KCl, 0.2% NP-40, 20% glycerol, 2 mM DTT) supplemented with 50 ng or 1 μg salmon sperm DNA (for pure protein and nuclear extract, respectively). 25 nM double-stranded probes were added, and reactions were incubated for 15 min at 37 °C before loading on native 6% PAGE gels at 150 V for 50 min. Gels were imaged using the Odyssey image analysis system (LiCor Biosciences). The double-stranded sequence of the *SKP2* probes used in EMSAs were Gccgaccagtcccgctcccgcggggggttgtgggtatctcgaaggcgggtaaagctgca (WT *SKP2* probe) and GccgaccagtcccgctcccgcggggggttgtgggtatctcgaaAAAAAAtaaagctgca (mutant *SKP2* probe). The WT probe was IRDye-800 fluorescence-labelled. In competition assays, unlabelled probes were included in binding reactions at 100-times the concentration of labelled probes. Analysis and quantification were performed using Image Studio Lite Ver 5.2.

### ChIP-qPCR

ChIP was performed as previously described^59^, using U2OS cells fixed in 1% formaldehyde for 15 min, ensuring sonication fragments between 100 and 500 bp, and using 5 μg anti-NUCKS1 antibody (ProteinTech 12023-2-AP) or 5 μg normal rabbit IgG (SantaCruz sc2027). Primers used for ChIP-qPCR are listed in Supplementary Table 4.

### Flow cytometry

For propidium iodide staining, trypsinised cells were fixed in cold 70% ethanol for 30 min on ice. Cells were then centrifuged at 250 g for 5 min and resuspended in PBS with 0.5 μg/ml RNAseA and 10 μg/ml propidium iodide, before incubation for 15 min at 37 °C. For EdU/PI staining, the Click-iT™ Plus EdU Alexa Fluor™ 488 Flow Cytometry Assay Kit was used, according to the manufacturer’s instructions.

The BD FACSCalibur™ (BD Biosciences) or CytoFLEX (Beckman Coulter) machines were used for sample acquisition. FlowJo v10.6.1 and ModFit LT 4.1.7 were used for analysis.

### Bioinformatics

Bioinformatics screens for putative transcriptional targets of NUCKS1, as outlined in Supplementary Fig. 1A, were performed using the SEEK database^47^ (http://seek.princeton.edu/). The SEEK database was used to generate lists of the 1000 genes correlating most positively with *NUCKS1* across 15 different cancer types, with three sample types per cancer (cancer tissue, tumour tissue, or cell line). This generated 45 lists of 1,000 genes, which were subsequently, independently, filtered through NUCKS1-interacting promoters in ChIP-Seq data_16_ (Supplementary Fig. 1A). The GO biological processes enrichment presented in Supplementary Fig. 1A was generated using Genecodis3^69^ (http://genecodis.cnb.csic.es). Fig. 1A and supplementary Fig. 1C and 1D were generated using data from GEPIA2^70^ (http://gepia2.cancer-pku.cn/#index). Fig. 6A and B were generated using data from UCSC XENA (https://xena.ucsc.edu/), using RSEM norm count values from GTEX/TCGA normal datasets, and TCGA tumour datasets^71^. Fig. 6C, D and F were generated using data from CBioPortal^72,73^ (https://www.cbioportal.org/).

### Statistical analyses

Statistical tests, indicated in figure legends, were performed using GraphPad Prism 8.

