## Supplementary data for "The NUCKS1-SKP2-p21/p27 axis controls S phase entry"

Hume et al.

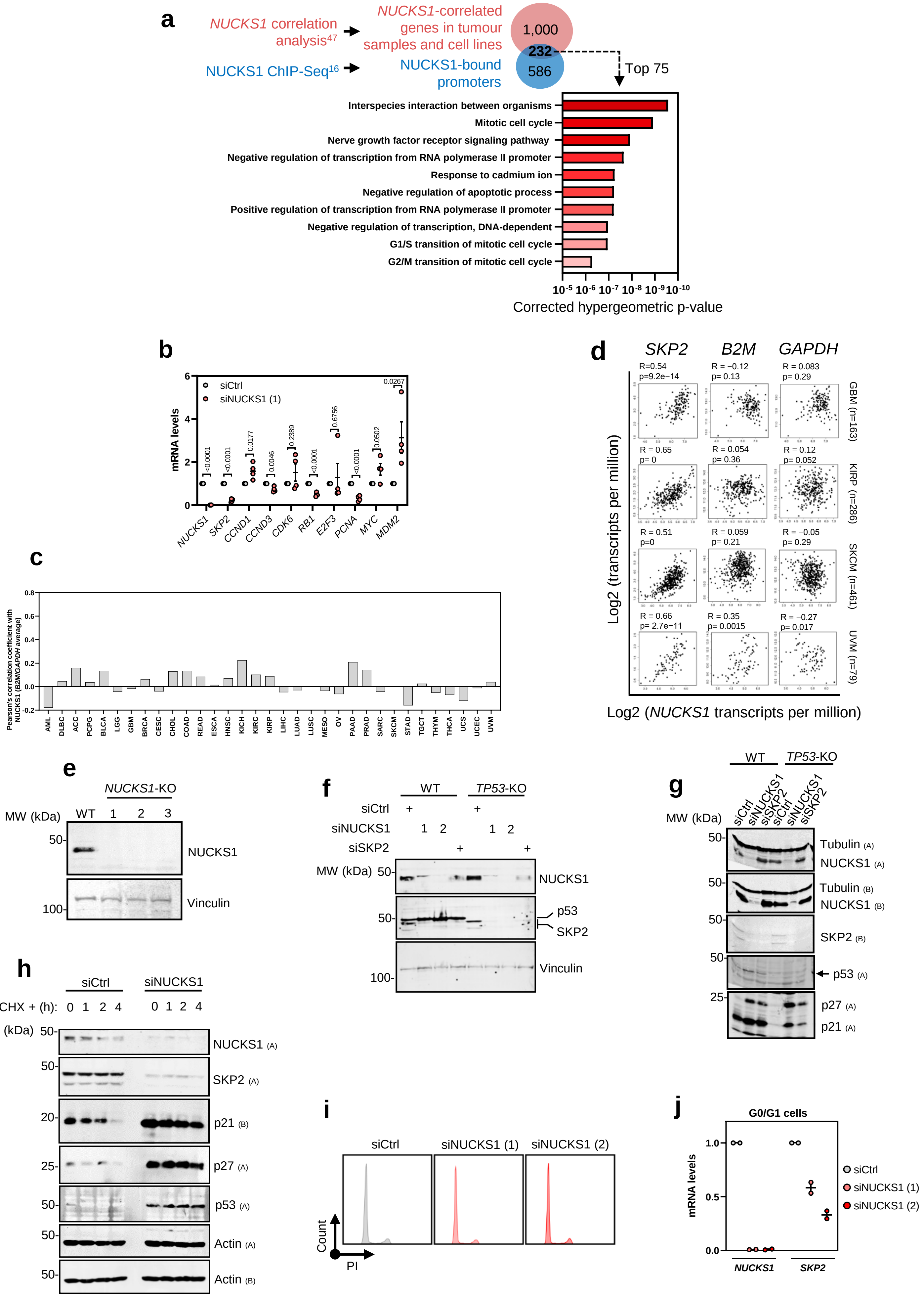
a
NUCKS1 correlation analysis47
NUCKS1 ChIP-Seq16
NUCKS1-correlated genes in tumour samples and cell lines
NUCKS1-bound promoters
1,000
232
586
Top 75
Interspecies interaction between organisms
Mitotic cell cycle
Nerve growth factor receptor signaling pathway
Negative regulation of transcription from RNA polymerase II promoter
Response to cadmium ion
Negative regulation of apoptotic process
Positive regulation of transcription from RNA polymerase II promoter
Negative regulation of transcription, DNA-dependent
G1/S transition of mitotic cell cycle
G2/M transition of mitotic cell cycle
10-5 10-6 10-7 10-8 10-9 10-10
Corrected hypergeometric p-value
b
mRNA levels
siCtrl
siNUCKS1 (1)
NUCKS1 SKP2 CCND1 CCND3 CDK6 RB1 E2F3 PCNA MYC MDM2
<0.0001 <0.0001 0.0177 0.0046 0.2389 <0.0001 0.6756 <0.0001 0.0502 0.0267
c
Pearson's correlation coefficient with NUCKS1 (B2M/GAPDH average)
AML DLBC ACC PCPG BLCA LGG GBM BRCA CESC CHOL COAD READ ESCA HNSC KICH KIRC KIRP LIHC LUAD LUSC MESO OV PAAD PRAD SARC SKCM STAD TGCT THYM THCA UCS UCEC UVM
d
SKP2 B2M GAPDH
R=0.54 p=9.2e−14
R = −0.12 p= 0.13
R = 0.083 p= 0.29
GBM (n=163)
R = 0.65 p= 0
R = 0.054 p= 0.36
R = 0.12 p= 0.052
KIRP (n=286)
R = 0.51 p=0
R = 0.059 p= 0.21
R = −0.05 p= 0.29
SKCM (n=461)
R = 0.66 p= 2.7e−11
R = 0.35 p= 0.0015
R = −0.27 p= 0.017
UVM (n=79)
Log2 (transcripts per million)
Log2 (NUCKS1 transcripts per million)
e
NUCKS1-KO
MW (kDa) WT 1 2 3
NUCKS1
Vinculin
f
WT TP53-KO
siCtrl
siNUCKS1 1 2
siSKP2
MW (kDa)
NUCKS1
p53
SKP2
Vinculin
g
WT TP53-KO
siCtrl siNUCKS1 siSKP2 siCtrl siNUCKS1 siSKP2
MW (kDa)
Tubulin (A)
NUCKS1 (A)
Tubulin (B)
NUCKS1 (B)
SKP2 (B)
p53 (A)
p27 (A)
p21 (A)
h
siCtrl siNUCKS1
CHX + (h): 0 1 2 4 0 1 2 4
MW (kDa)
NUCKS1 (A)
SKP2 (A)
p21 (B)
p27 (A)
p53 (A)
Actin (A)
Actin (B)
i
siCtrl siNUCKS1 (1) siNUCKS1 (2)
Count
PI
j
G0/G1 cells
mRNA levels
siCtrl
siNUCKS1 (1)
siNUCKS1 (2)
NUCKS1 SKP2

**Supplementary Figure 1: NUCKS1 transcriptionally controls the SKP2-p21/p27 axis (related to Figure 1)**

A) Schematic of approach to identify NUCKS1's transcriptional targets, and the GO biological processes enrichment of the top hits, using NUCKS1 ChIP-Seq[16], SEEK[47], and GeneCodis[69].
B) RT-qPCR in WT RPE1-hTERT cells treated with control or NUCKS1 siRNA. Two-tailed Student's T test.
C) Pearson's correlation (two-tailed) of *B2M*/*GAPDH* (used as control, housekeeping genes) and *NUCKS1* mRNAs in cancer, using data from Gepia2[70].
D) Pearson's correlation (two-tailed) of *NUCKS1* and *SKP2*/*B2M*/*GAPDH* mRNAs in cancer, made using Gepia2[70]. GBM = Glioblastoma, KIRP = Kidney renal papillary cell carcinoma, SKCM = Skin Cutaneous Melanoma, UVM = Uveal melanoma.
E) Western blot validating CRISPR/Cas9-mediated *NUCKS1* deletion in U2OS cells.
F) Western blot in control- or NUCKS1-depleted WT or *TP53*-KO RPE1-hTERT cells.
G) Western blot in control, NUCKS1-, or SKP2-depleted WT and *TP53*-KO RPE1-hTERT cells.
H) Western blot from WT RPE1-hTERT cells treated with control or NUCKS1 siRNA followed by treatment with 10 µg/ml cycloheximide (CHX) for the indicated periods of time.
I) Cell cycle profiles of NBE1-hTERT cells synchronised to G0/G1 by contact inhibition for 72 h followed by control or NUCKS1 depletion for 72 further hours.
J) RT-qPCR of NBE1-hTERT cells treated as in I.

In E, F, G and H, data are representative of 3 (E, F, G) or 2 (H) independent experiments. In B and J, data are presented as mean +/- SEM of 4 or 2 independent experiments, respectively.

MW: molecular weight, kDa: kilodaltons, PI: propidium iodide. Source data are provided as a source data file.

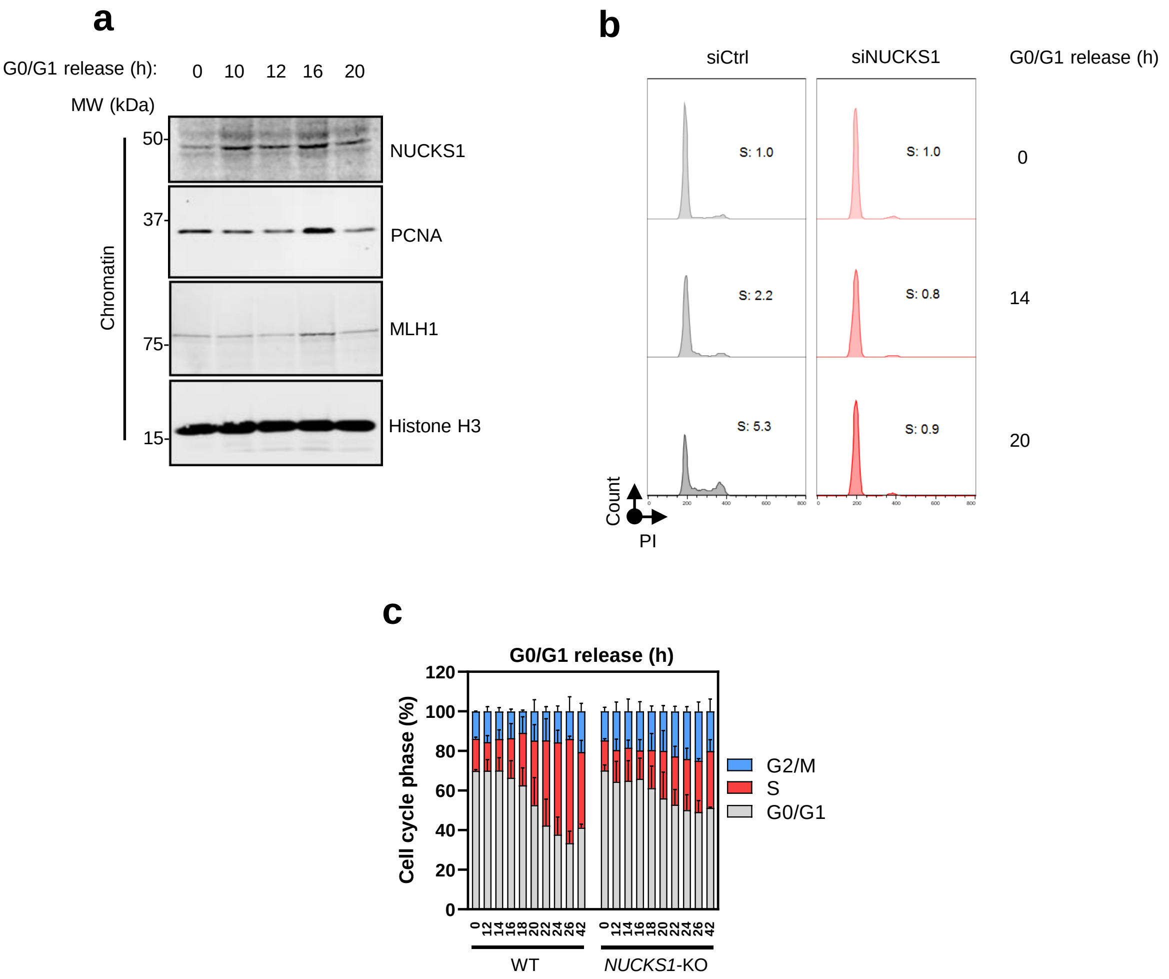


**Supplementary Figure 2: NUCKS1 levels and chromatin-binding are induced in late G1 to promote *SKP2* expression and G1/S progression (related to figure 2)**

A) Western blot in NBE1-hTERT cells synchronised to G0/G1 by contact inhibition for 72 h (t=0) and released to enter the cell cycle by re-plating at low density.

B) PI profiles of NBE1-hTERT cells synchronised to G0/G1 by contact inhibition for 72 h (*t*=0), treated with control or NUCKS1 siRNA for a further 72 h, and released to enter the cell cycle by re-plating at low density.

C) Cell cycle quantification of WT or *NUCKS1*-KO U2OS cells synchronised to G0/G1 by treating with 40 µM lovastatin for 48 h (*t*=0) followed by 4 mM mevalonate to release cells into S phase.

In A and B, data are representative of 2 or 3 independent experiments, respectively. In C, data are presented as mean +/- SEM of 2 independent experiments.

MW: molecular weight, kDa: kilodaltons, PI: propidium iodide. Source data are provided as a source data file.

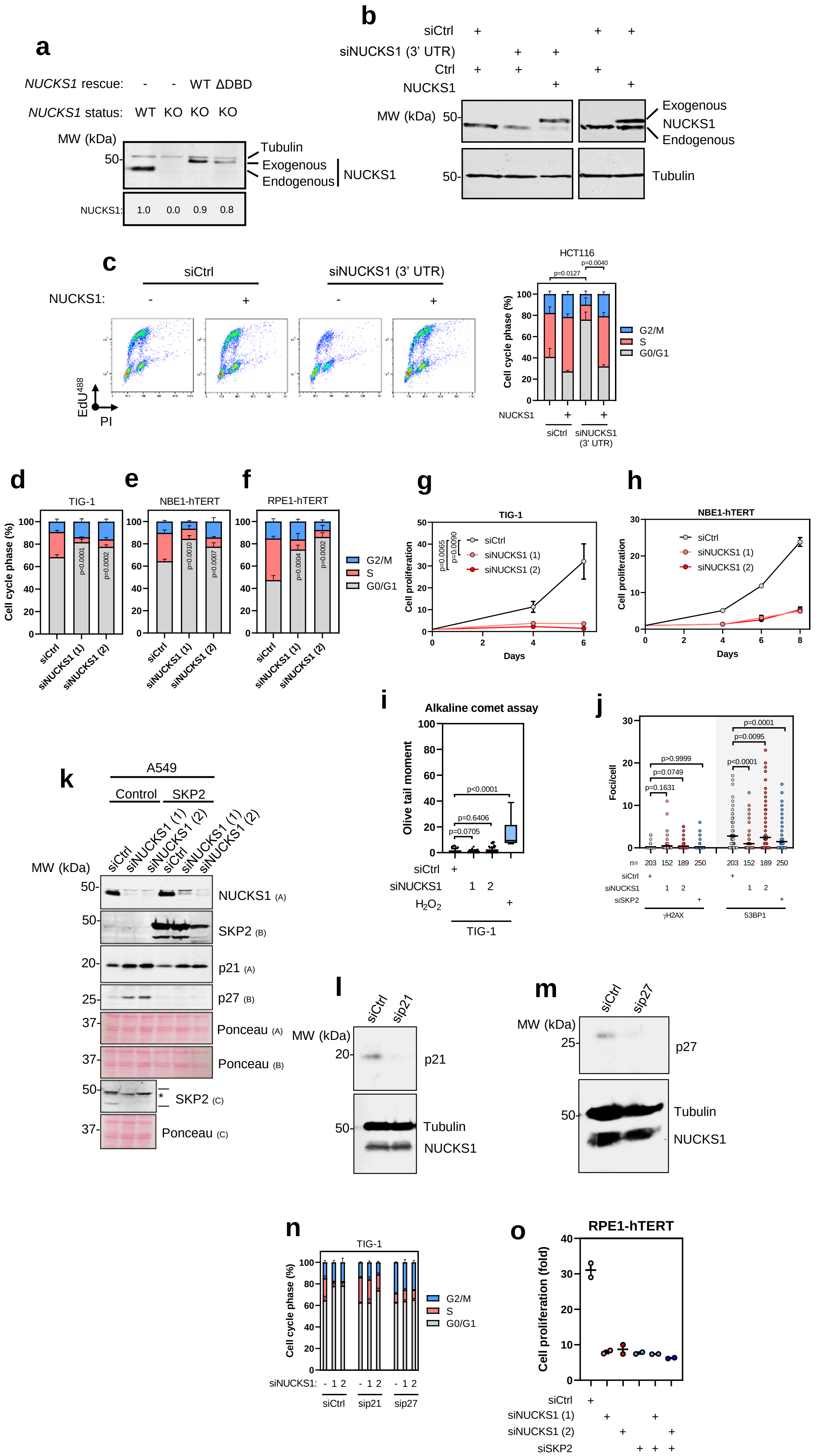
a
NUCKS1 rescue: - - WT ΔDBD
NUCKS1 status: WT KO KO KO
MW (kDa)
50-
Tubulin
Exogenous
Endogenous
NUCKS1
NUCKS1: 1.0 0.0 0.9 0.8
b
siCtrl
siNUCKS1 (3' UTR)
Ctrl
NUCKS1
MW (kDa) 50-
Exogenous
NUCKS1
Endogenous
50-
Tubulin
c
siCtrl
siNUCKS1 (3' UTR)
NUCKS1: - + - +
EdU488
PI
HCT116
p=0.0127
p=0.0040
Cell cycle phase (%)
G2/M
S
G0/G1
NUCKS1
siCtrl
siNUCKS1 (3' UTR)
d
TIG-1
p<0.0001
p=0.0002
siCtrl
siNUCKS1 (1)
siNUCKS1 (2)
e
NBE1-hTERT
p=0.0010
p=0.0007
f
RPE1-hTERT
p=0.0004
p=0.0002
G2/M
S
G0/G1
g
TIG-1
p=0.0065
p=0.0090
siCtrl
siNUCKS1 (1)
siNUCKS1 (2)
Cell proliferation
Days
h
NBE1-hTERT
siCtrl
siNUCKS1 (1)
siNUCKS1 (2)
Cell proliferation
Days
i
Alkaline comet assay
Olive tail moment
p<0.0001
p=0.6406
p=0.0705
siCtrl +
siNUCKS1 1 2
H2O2 +
TIG-1
j
Foci/cell
p>0.9999
p=0.0749
p=0.1631
p=0.0001
p=0.0095
p<0.0001
n= 203 152 189 250 203 152 189 250
siCtrl
siNUCKS1
siSKP2
γH2AX
53BP1
k
A549
Control SKP2
siCtrl siNUCKS1 (1) siNUCKS1 (2) siCtrl siNUCKS1 (1) siNUCKS1 (2)
MW (kDa)
NUCKS1 (A)
SKP2 (B)
p21 (A)
p27 (B)
Ponceau (A)
Ponceau (B)
SKP2 (C)
Ponceau (C)
l
siCtrl sip21
MW (kDa)
p21
Tubulin
NUCKS1
m
siCtrl sip27
MW (kDa)
p27
Tubulin
NUCKS1
n
TIG-1
Cell cycle phase (%)
G2/M
S
G0/G1
siNUCKS1: - 1 2 - 1 2 - 1 2
siCtrl sip21 sip27
o
RPE1-hTERT
Cell proliferation (fold)
siCtrl
siNUCKS1 (1)
siNUCKS1 (2)
siSKP2

**Supplementary Figure 3: NUCKS1 controls S phase entry through the SKP2-p21/p27 axis (related to Figure 3)**

A) Western blot in U2OS cells expressing indicated variants of NUCKS1.

B) Western blot validating NUCKS1 depletion and overexpression in HCT116 cells.

C) EdU/PI cell cycle profiles of HCT116 cells (left) and corresponding quantifications (right). Ordinary one-way ANOVA with Sidak multiple comparisons test on S phase population.

D) Cell cycle quantification of TIG-1 cells. For D-F, ordinary one-way ANOVAs with Dunnett multiple comparisons test on the S phase populations were performed.

E) Cell cycle quantification of NBE1-hTERT cells.

F) Cell cycle quantification of RPE1-hTERT cells.

G) Proliferation assay in TIG-1 cells. One-way ANOVA with Dunnett multiple comparisons test at day 6.

H) Proliferation assay in NBE1-hTERT cells.

I) DNA damage measured using the alkaline comet assay in TIG-1 cells. Treatment for 10 min with 50 µM $H_2O_2$ was used as a positive control. N=103, 98, 108, 9, for siCtrl, siNUCKS1 (1, 2), $H_2O_2$, respectively. Box plots show median values along with 25/75% (box) and 10/90% (whiskers) and outliers. Kruskal-Wallis with Dunn's multiple comparisons test. Representative of 2 independent experiments.

J) DNA damage measured using γH2AX or 53BP1 immunofluorescence in RPE1-hTERT cells. Mean of n=203, 152, 189, and 250 cells +/- SEM for control, NUCKS1 (sequence 1/2) and SKP2 depletions, respectively. Kruskal-Wallis with Dunn's multiple comparisons test. Representative of one experiment.

K) Western blot in A549 cells. Asterisk indicates a non-specific band.

L) Western blot validating p21 depletion by siRNA

M) Western blot validating p27 depletion by siRNA.

N) Cell cycle quantification of TIG-1 cells.

O) Fold proliferation increase at day 4 after siRNA in RPE1-hTERT cells.

In A, B, C (left), K, L, and M, data are representative of 2 (A, K, M), 1 (B) or 3 (C, L) independent experiments. In C (right), D, E, F, G, H, N and O, data are presented as mean +/- SEM of 3 (C, D, G), 4 (E), 6 (F), or 2 (H, N, O) experiments.

MW: molecular weight, kDa: kilodaltons, PI: propidium iodide. Source data are provided as a source data file.

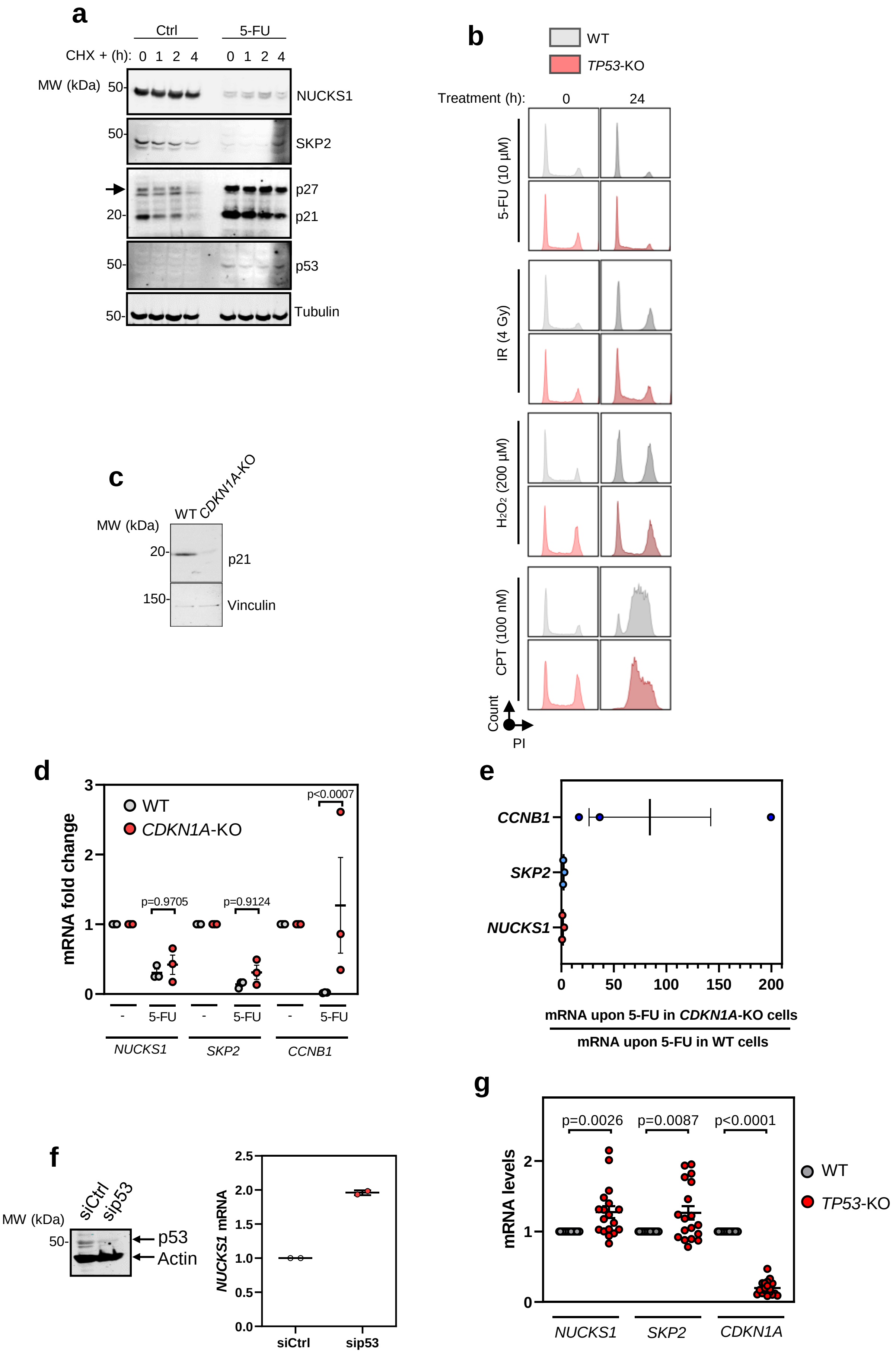
a
Ctrl
5-FU
CHX + (h): 0 1 2 4 0 1 2 4
MW (kDa)
50-
NUCKS1
50-
SKP2
p27
20-
p21
50-
p53
50-
Tubulin
b
WT
TP53-KO
Treatment (h): 0 24
5-FU (10 µM)
IR (4 Gy)
H2O2 (200 µM)
CPT (100 nM)
Count
PI
c
WT
CDKN1A-KO
MW (kDa)
20-
p21
150-
Vinculin
d
mRNA fold change
WT
CDKN1A-KO
p=0.9705
p=0.9124
p<0.0007
- 5-FU - 5-FU - 5-FU
NUCKS1
SKP2
CCNB1
e
CCNB1
SKP2
NUCKS1
0 50 100 150 200
mRNA upon 5-FU in CDKN1A-KO cells
mRNA upon 5-FU in WT cells
f
siCtrl
sip53
MW (kDa)
50-
p53
Actin
NUCKS1 mRNA
siCtrl
sip53
g
mRNA levels
p=0.0026
p=0.0087
p<0.0001
WT
TP53-KO
NUCKS1
SKP2
CDKN1A

**Supplementary Figure 4: DNA damage inhibits the NUCKS1-SKP2 axis through p53-mediated transcriptional repression (related to Figure 5)**

A) Western blot in control or 5-FU (10 µM, 48 h)-treated WT RPE1-hTERT cells chased with cycloheximide (10 µg/ml) for the indicated periods of time.
B) Cell cycle profiles of WT and *TP53*-KO RPE1-hTERT cells treated with 10 µM 5-FU, 4 Gy IR, 200 µM $H_2O_2$, or 100 nM CPT for the indicated periods of time.
C) Western blot validation of *CDKN1A*-KO RPE1-hTERT cells.
D) RT-qPCR in RPE1-hTERT WT and *CDKN1A*-KO cells treated with 10 µM 5-FU for 24 h. One-way ANOVA with Sidak's test for multiple comparisons.
E) Data from D: mRNA levels in 5-FU-treated *CDKN1A*-knockout cells were divided by mRNA levels in 5-FU-treated WT cells.
F) RT-qPCR after p53 siRNA in TIG-1 cells.
G) RT-qPCR in untreated RPE1-hTERT WT and *TP53*-KO cells. Two-tailed Student's t test.

In A, B, C, and F (left), data are representative of 2 (A, F), 3 (B), or 1 (C) independent experiments. In D, E, F (right), and G, data are presented as mean +/- SEM of 3 (D, E), 2 (F), or 18 (G) independent experiments.

MW: molecular weight, kDa: kilodaltons, PI: propidium iodide. Source data are provided as a source data file.

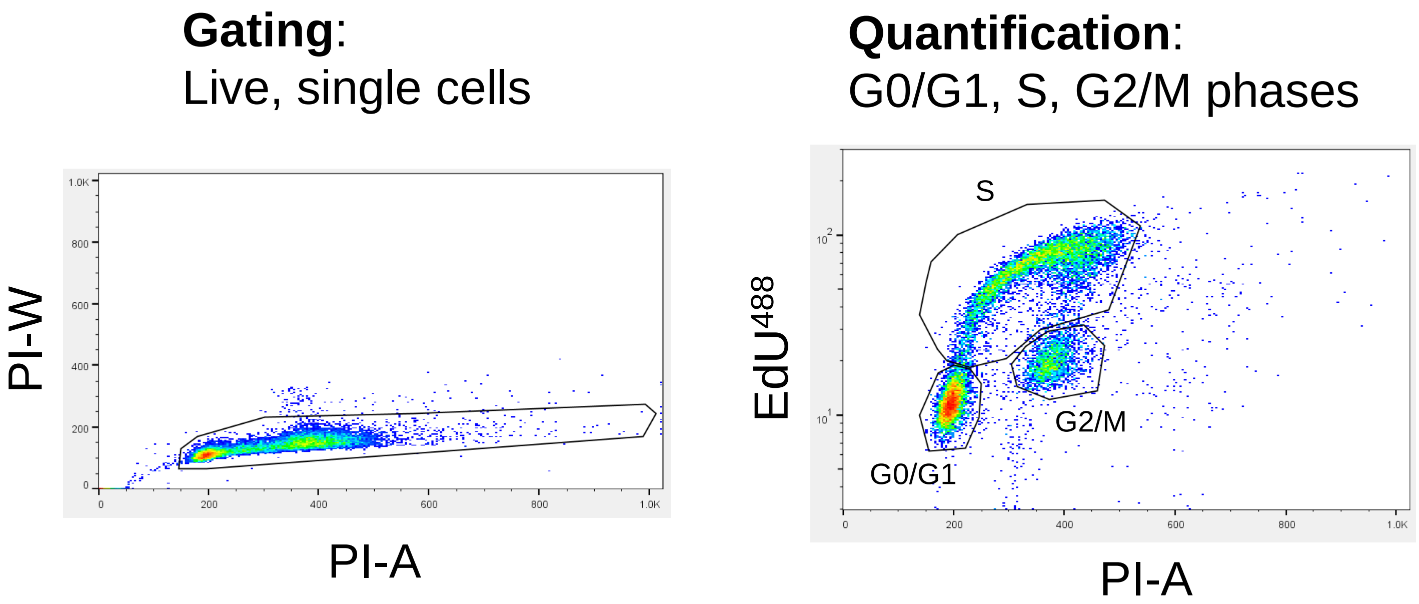


**Supplementary Figure 5: Gating strategy for FACS**

Gating strategy used for flow cytometry. The gate on the left was used to isolate live, single cells (used for analyses of both PI and EdU/PI), and the gate on the right was subsequently used to quantify EdU/PI profiles, based on the percentages of cells in G0/G1, S, and G2/M phases of the cell cycle.

| Supplementary table 1 | | |
|---|---|---|
| **Cell line** | **Source** | **Code** |
| TIG-1, primary embryonic fibroblasts | Coriell Institute Cell Repository | AG06173 |
| NBE1-hTERT, hTERT-immortalised normal bronchial epithelial cells | Prof. Anderson Ryan[74] | N/A |
| U2OS, Osteosarcoma | American Type Culture Collection (ATCC) | ATCC® HTB-96™ |
| U2OS *NUCKS1*-KO | This study | N/A |
| RPE1-hTERT WT, hTERT-immortalised retinal pigment epithelial cells | ATCC | ATCC® CRL-4000™ |
| RPE1-hTERT *TP53*-KO | Dr. Ross Chapman | N/A |
| RPE1-hTERT *CDKN1A*-KO | Prof. Iain Cheeseman[75] | N/A |
| RKO, colorectal cancer | ATCC | ATCC® CRL-2577™ |
| HCT116, colorectal cancer | ATCC | ATCC® CCL-247™ |
| HCT116 *TP53*-KO | Prof. Vogelstein[77] | N/A |
| CACO2, colorectal cancer | ATCC | ATCC® HTB-37™ |
| HT29, colorectal cancer | ATCC | ATCC® HTB-38™ |
| Doxycycline-inducible SKP2 A549, lung adenocarcinoma | Prof. Shyamala Maheswaran[76] | N/A |
| DLD1, colorectal cancer | ATCC | ATCC® CCL-221™ |
| Sf9 cells | ThermoFisher Scientific | 11496015 |

**Supplementary Table 1: Cell lines used in the study.**

| Supplementary table 2 | | |
|---|---|---|
| **Recombinant DNA** | **Source** | **Code** |
| SKP2 | Dr. Vincenzo D'Angiolella | N/A |
| p53 WT | Prof. Eric O'Neill | N/A |
| p53 R175H | Prof. Eric O'Neill | N/A |
| p53 R248W | Prof. Eric O'Neill | N/A |
| p53 R273H | Prof. Eric O'Neill | N/A |
| *NUCKS1* CRISPR/Cas9 KO plasmid | SantaCruz | sc-413018 |
| *NUCKS1* HDR plasmid | SantaCruz | sc-413018-HDR |
| NUCKS1 WT | Abgent (Generon) | DC00385 |
| NUCKS1 DNA-binding mutant (GRP to AAA) | This study | N/A |
| pDEST53-NUCKS1 | Thermofisher Scientific | 12288015 |

**Supplementary Table 2: Recombinant DNA used in the study.**

| Supplementary table 3 | | | |
|---|---|---|---|
| **Antibody** | **Source** | **Code** | **Dilution** |
| NUCKS1 | Prof. Anne Østvold[25] | N/A | 1:2000 |
| NUCKS1 | ProteinTech | 12023-2-AP | 1:1000 |
| SKP2 | ThermoFisher Scientific | 32-3300 | 1:1000 |
| p21 | Cell signalling | #2947 | 1:2000 |
| p27 | Cell signalling | #2552 | 1:500 |
| p53 | Santa Cruz | sc-126 | 1:500 |
| PCNA | Santa Cruz | sc-56 | 1:1000 |
| MLH1 | Abcam | ab92312 | 1:2000 |
| Beta-Actin | Abcam | ab6276 | 1:10000 |
| Alpha-Tubulin | Sigma-Aldrich | T6199 | 1:5000 |
| Alpha-Tubulin | Abcam | Ab4074 | 1:5000 |
| Vinculin | Santa Cruz | sc-73614 | 1:1000 |
| Histone H3 | Abcam | ab201456 | 1:2000 |
| Cyclin A2 | Abcam | ab181591 | 1:2000 |
| IRDye® 800CW Goat anti-Rabbit IgG (H + L) | Li-Cor | 925-32211 | 1:10000 |
| IRDye® 680RD Goat anti-Mouse IgG (H + L) | Li-Cor | 925-68070 | 1:10000 |
| IRDye® 680RD Goat anti-Rabbit IgG Secondary Antibody | Li-Cor | 925-68071 | 1:10000 |
| IRDye® 800CW Goat anti-Mouse IgG Secondary Antibody | Li-Cor | 926-32210 | 1:10000 |
| Normal rabbit IgG | Santa Cruz | sc-2027 | |
| Normal mouse IgG | Santa Cruz | sc-2025 | |
| γH2AX | Millipore | 05-636 | 1:250 |
| 53BP1 | Santa Cruz | sc-22760 | 1:250 |
| Alexa Fluor 488, Goat Anti-Rabbit IgG (H + L) | Thermo Fisher Scientific | A-11034 | 1:500 |
| Alexa Fluor 555, Goat Anti-Mouse IgG (H + L) | Thermo Fisher Scientific | A-32727 | 1:500 |

**Supplementary Table 3: Antibodies used in the study.**

| Supplementary table 4 | |
|---|---|
| **Primers for RT-qPCR** | **Primers for ChIP-qPCR** |
| *NUCKS1*-F; GCCCGAATCTCTTCCATAATC | *SKP2* promoter F: -9806 - -9695 TCTGAGATAGGCTGAGAGGAAA |
| *NUCKS1*-R; TTCTCGACTCGGTCCTGTTT | *SKP2* promoter R: -9806 - -9695 CTTCAGTAGACGGTAGTGAGAAAC |
| *SKP2*-F; GGAAGGGAGTCCCATGAAA | *SKP2* promoter F: -1448 - -1298 CAATCTTCAGGGAAAGGACGTAG |
| *SKP2*-R; GCTGAAGAGCAAAGGGAGTG | *SKP2* promoter R: -1448 - -1298 CTAGGCTAAGCCGTTCATCAAA |
| *CDKN1A*–F; CATGGGTTCTGACGGACATC | *SKP2* promoter F: -855 - -673 CAGCAGAGATAGGCTGAAAG |
| *CDKN1A*–R; TGCCGAAGTCAGTTCCTTGT | *SKP2* promoter R: -855 - -673 GCCTTGACAGGTCTCATAAA |
| *CDKN1B*–F; TTCATCAAGCAGTGATGTATCTGA | *SKP2* promoter F: -417 - -266 GCTACTGTCACCACTCAGCA |
| *CDKN1B*–R; AAGAAGCCTGGCCTCAGAAG | *SKP2* promoter R: -417 - -266 GATCGGACGGTGAGCCTAAG |
| *B2M-F;* ATGTCTCGCTCCGTGGCCTTA | *SKP2* promoter F: -346 - -197 CTCTCTTTCCGCCTGCTCTC |
| *B2M-R;* ATCTTGGGCTGTGACAAAGTC | *SKP2* promoter R: -346 - -197 GTGGATCCCAGAACCTGGAC |
| *GAPDH*-F; AGCCACATCGCTCAGACAC | *SKP2* promoter F: -25 - +148 GTATCTCGAAGGCGGGTAAAG |
| *GAPDH*-R; GCCCAATACGACCAAATCC | *SKP2* promoter R: -25 - +148 TAAGCCTAGCAACGTTCCATC |
| *TBP*-F; CGGTTTGCTGCGGTAATCAT | *SKP2* promoter F: +420 - +575 GTGAATGGATGGATGCCTTAG |
| *TBP*-R; TTTCTTGCTGCCAGTCTGGAC | *SKP2* promoter R: +420 - +575 GCAGAAGGCCTCAAGAATTT |
| *CCNB1*-F; CGGGAAGTCACTGGAAACAT | *SKP2* promoter F: +843 - +1001 TGGGATTCCAGCAAGACTTC |
| *CCNB1*-R; AAACATGGCAGTGACACCAA | *SKP2* promoter R: +843 - +1001 GTCACTCCCTTTGCTCTTCA |
| *BBC3*-F; GTAAGGGCAGGAGTCCCAT | *SKP2* promoter F: +1455 - +1592 TATCAGTGGACTGCGAAGTGA |
| *BBC3*-R; GACGACCTCAACGCACAGTA | *SKP2* promoter R: +1455 - +1592 CGGGTTGGGTGATTAAACAGATAG |
| *MDM2*-F; GCAGTGAATCTACAGGGACGC | *SKP2* promoter F: +10467 - +10629 CTCTCCCTTGCCCTGTTTAAT |
| *MDM2*-R; ATCCTGATCCAACCAATCACC | *SKP2* promoter R: +10467 - +10629 AGGTGTTGGGAAGGAGTAGTA |
| *TP53*-F; TGTTTCCTGACTCAGAGGGG | |
| *TP53*-R; GAGCGTGCTTTCCACGAC | |
| *RB1*-F; TCAGTTGGTCCTTCTCGGTC | |
| *RB1*-R; TGTGAACATCGAATCATGGAA | |
| *CDK6*-F; CACTCCAGGCTCTGGAACTT | |
| *CDK6*-R; TGGAGACCTTCGAGCACC | |
| *CCND1*-F; AGTTGTTGGGGCTCCTCAG | |
| *CCND1*-R; AGACCTTCGTTGCCCTCTGT | |
| *MYC*-F; CACCGAGTCGTAGTCGAGGT | |
| *MYC*-R; TTTCGGGTAGTGGAAAACCA | |
| *PCNA*-F; AAGAGAGTGGAGTGGCTTTTG | |
| *PCNA*-R; TGTCGATAAAGAGGAGGAAGC | |
| *MDM2*-F; GCAGTGAATCTACAGGGACGC | |
| *MDM2*-R; ATCCTGATCCAACCAATCACC | |
| *E2F3*-F; AAGTGCCTGACTCAATAGAGAGCC | |
| *E2F3*-R; AGTCTCTTCTGGACATAAGTAAACCTCA | |
| *CCND3*-F; TTGAGCTTCCCTAGGACCAG | |
| *CCND3*-R; TGACCATCGAAAAACTGTGC | |

**Supplementary Table 4: Primers used in the study.**
